# Long-read transcript sequencing identifies differential isoform expression in the entorhinal cortex in a transgenic model of tau pathology

**DOI:** 10.1101/2023.09.20.558220

**Authors:** Szi Kay Leung, Aaron R Jeffries, Isabel Castanho, Rosemary A Bamford, Karen Moore, Emma L Dempster, Jonathan T Brown, Zeshan Ahmed, Paul O’Neill, Eilis Hannon, Jonathan Mill

## Abstract

Increasing evidence suggests that alternative splicing plays an important role in Alzheimer’s disease (AD), a devastating neurodegenerative disorder involving the intracellular aggregation of hyperphosphorylated tau. We used long-read cDNA sequencing to profile transcript diversity in the entorhinal cortex of wild-type (WT) and transgenic (TG) mice harboring a mutant form of human tau. Whole transcriptome profiling showed that previously reported gene-level expression differences between WT and TG mice reflect changes in the abundance of specific transcripts. Ultradeep targeted long-read cDNA sequencing of genes implicated in AD revealed hundreds of novel isoforms and identified specific transcripts associated with the development of tau pathology. Our results highlight the importance of differential transcript usage, even in the absence of gene-level expression alterations, as a mechanism underpinning gene regulation in the development of neuropathology. Our transcript annotations and a novel informatics pipeline for the analysis of long-read transcript sequencing data are provided as a resource to the community.

## Introduction

Alzheimer’s disease (AD) is a devastating neurodegenerative disorder that is clinically characterized by progressive memory loss, cognitive decline, and behavioral impairment^1^. These symptoms result from brain atrophy, synaptic loss and progressive neuropathology involving the extracellular accumulation of β-amyloid (Aβ) proteins in amyloid plaques and the intracellular aggregation of hyperphosphorylated tau into neurofibrillary tangles^2^. Despite recent success in identifying genetic risk factors for AD, the mechanisms driving disease progression remain unclear.

Alternative splicing (AS) is a post-transcriptional regulatory mechanism that generates multiple RNA isoforms from a single transcribed messenger RNA (mRNA) precursor. By dramatically increasing transcriptional diversity from the coding genome, AS represents an important regulator of gene expression. Transcriptional dysregulation, and in particular aberrant splicing, has been shown to play a key role in the development and pathogenesis of AD^3,4^ with isoform expression differences identified in post-mortem brain tissue from AD patients^3,4^ and mouse models with AD pathology^5,6^. Previous studies investigating AS have been limited by the use of short-read RNA sequencing (RNA-Seq) approaches, which cannot span full-length transcripts and unequivocally detect or quantify specific isoforms^7^. We recently demonstrated the power of long-read sequencing for the characterization of transcript diversity and quantification of alternatively-spliced isoforms in the mouse cortex^8^.

In this study, we utilize two complementary long-read sequencing approaches – Pacific Biosciences (PacBio) isoform sequencing (Iso-Seq) and Oxford Nanopore Technologies (ONT) nanopore cDNA sequencing – to examine isoform diversity and AS in a well-characterized transgenic model of tau pathology (rTg4510)^9^. These mice over-express a human mutant (P301L) form of the microtubule-associated protein tau (MAPT) and are characterized by widespread transcriptional dysregulation paralleling the accumulation of tau pathology in the cortex^9^. The spread of tau pathology in rTg4510 mice^5^ closely recapitulates the Braak stages in AD^10^, with appearance of pre-tangles from as early as 3 months followed by synaptic and neuronal loss by 9 months. First, we performed whole transcriptome long-read sequencing of the entorhinal cortex (EC) – a brain region defined by early neuropathology in AD^10^ – profiling tissue from both wild-type (WT) and transgenic (TG) mice at two time-points (2 and 8 months) that are characterized by early and late stages of tauopathy^9^. Second, we performed ultra-deep targeted long-read sequencing of transcripts expressed from 20 genes previously implicated in AD and other neurodegenerative disorders (**Table 1**) using tissue from WT and TG mice at four time-points (2, 4, 6 and 8 months). Finally, we developed a novel informatics tool, *FICLE* (<u>F</u>ull Isoform <u>C</u>haracterisation from <u>L</u>ong-read sequencing <u>E</u>xperiments), that facilitates the accurate assessment of AS events and the visualization of isoforms based on splicing patterns, leveraging the high sequencing coverage of targeted long-read transcriptome datasets. Our transcript annotations and data analysis pipeline are available as a resource to the research community (see **Data Availability**). Our results confirm the importance of AS in the mouse cortex and provide evidence of differential transcript usage, even in the absence of gene-level expression alterations, associated with the development of tau pathology in TG mice.

**Table 1:** Diversity of transcripts expressed from the 20 genes characterized using ultra-deep targeted long-read sequencing (ONT nanopore and PacBio Iso-Seq). Shown for each gene are the number of known and novel isoforms, further classified by coding potential, the number of isoforms characterized by alternative splicing events (A5A3 - alternative 5’ and 3’ splice site, ES - exon skipping, IR - intron retention) and the total number of these events across all associated isoforms.

| Target genes | Number of isoforms (merged ONT and PacBio dataset) |  |  |  |  | Number of isoforms with AS events<br>(total number of AS events) |  |  |
| --- | --- | --- | --- | --- | --- | --- | --- | --- |
|  | Classification |  |  | Coding potential |  | A5A3 | ES | IR |
|  | All | Known | Novel | Coding | Non-coding |  |  |  |
| *Abca1*<sup>28</sup> | 44 | 34 | 10 | 43 | 1 | 30 (45) | 3 (3) | 0 (0) |
| *Abca7*<sup>29</sup> | 82 | 36 | 46 | 81 | 1 | 56 (80) | 26 (47) | 1 (1) |
| *Ank1*<sup>30</sup> | 54 | 44 | 10 | 44 | 10 | 31 (40) | 37 (116) | 0 (0) |
| *Apoe* | 1408 | 13 | 1395 | 1348 | 60 | 1380 (2650) | 1303 (4316) | 11 (11) |
| *App*<sup>31</sup> | 1119 | 147 | 972 | 1074 | 45 | 909 (1436) | 1037 (5889) | 0 (0) |
| *Bin1*<sup>20</sup> | 624 | 52 | 572 | 605 | 19 | 476 (700) | 571 (2821) | 0 (0) |
| *Cd33*<sup>32</sup> | 54 | 16 | 38 | 51 | 3 | 50 (91) | 52 (83) | 9 (10) |
| *Clu*<sup>20</sup> | 964 | 28 | 936 | 937 | 27 | 915 (1758) | 705 (3894) | 1 (1) |
| *Fus*<sup>20</sup> | 312 | 74 | 238 | 282 | 30 | 234 (292) | 281 (1428) | 36 (36) |
| *Fyn*<sup>33</sup> | 96 | 45 | 51 | 93 | 3 | 72 (98) | 79 (286) | 0 (0) |
| *Mapt*<sup>34</sup> | 111 | 22 | 89 | 72 | 39 | 94 (132) | 88 (404) | 0 (0) |
| *Picalm*<sup>35</sup> | 323 | 140 | 183 | 232 | 91 | 176 (206) | 310 (839) | 0 (0) |
| *Ptk2b*<sup>36</sup> | 672 | 85 | 587 | 667 | 5 | 513 (811) | 416 (1287) | 1 (1) |
| *Rhbdf2*<sup>37</sup> | 18 | 12 | 6 | 18 | 0 | 9 (10) | 2 (2) | 0 (0) |
| *Snca*<sup>38</sup> | 725 | 14 | 711 | 281 | 444 | 690 (1382) | 712 (1732) | 1 (1) |
| *Sorl1*<sup>39</sup> | 213 | 137 | 76 | 199 | 14 | 197 (280) | 94 (94) | 0 (0) |
| *Tardbp*<sup>40</sup> | 191 | 79 | 112 | 150 | 41 | 113 (157) | 188 (907) | 69 (69) |
| *Trem2*<sup>41</sup> | 97 | 13 | 84 | 80 | 17 | 86 (152) | 90 (129) | 5 (5) |
| *Trpa1*<sup>42</sup> | 4 | 4 | 0 | 4 | 0 | 1 (1) | 0 (0) | 0 (0) |
| *Vgf*<sup>43</sup> | 51 | 5 | 46 | 44 | 7 | 50 (75) | 37 (38) | 0 (0) |

## Results

### Whole transcriptome long-read sequencing identifies specific transcripts of Gfap associated with the progression of tau pathology

We previously used short-read RNA-Seq to identify dramatic changes in EC gene expression associated with the development of tau pathology in TG mice carrying a human mutant form of the MAPT gene^5^. In the current study, we used long-read cDNA sequencing to extend these analyses and profile AS and differential transcript expression in the same mice. Our first analyses used PacBio Iso-Seq to generate whole transcriptome long-read sequencing data from a subset of WT and TG mice (total n = 12, 3 WT and 3 TG at ages 2 and 8 months) (**Figure 1A**, **Supplementary Table 1**) with raw reads processed using a customized Iso-Seq bioinformatics pipeline (see **Methods**)^8^. Across samples, we detected an average of 10,950 (SD = 615) genes and 38,930 (SD = 2,016) transcripts with a mean length of 2.7kb (SD = 1.6kb) and 9.88 exons (SD = 7.56). There were no differences in gene or transcript characteristics between genotype (WT vs TG mice) or age groups (2 vs 8 months) (**Supplementary Figure 1**, **Supplementary Table 2**). As expected, human-specific *MAPT* transcripts were only detected in full-length reads from TG mice, confirming the stable activation of the human *MAPT* transgene (**Supplementary Figure 2**). Raw data and annotated transcripts from both WT and TG mice are available as a downloadable resource (see **Data Availability**).

**Figure 1:**
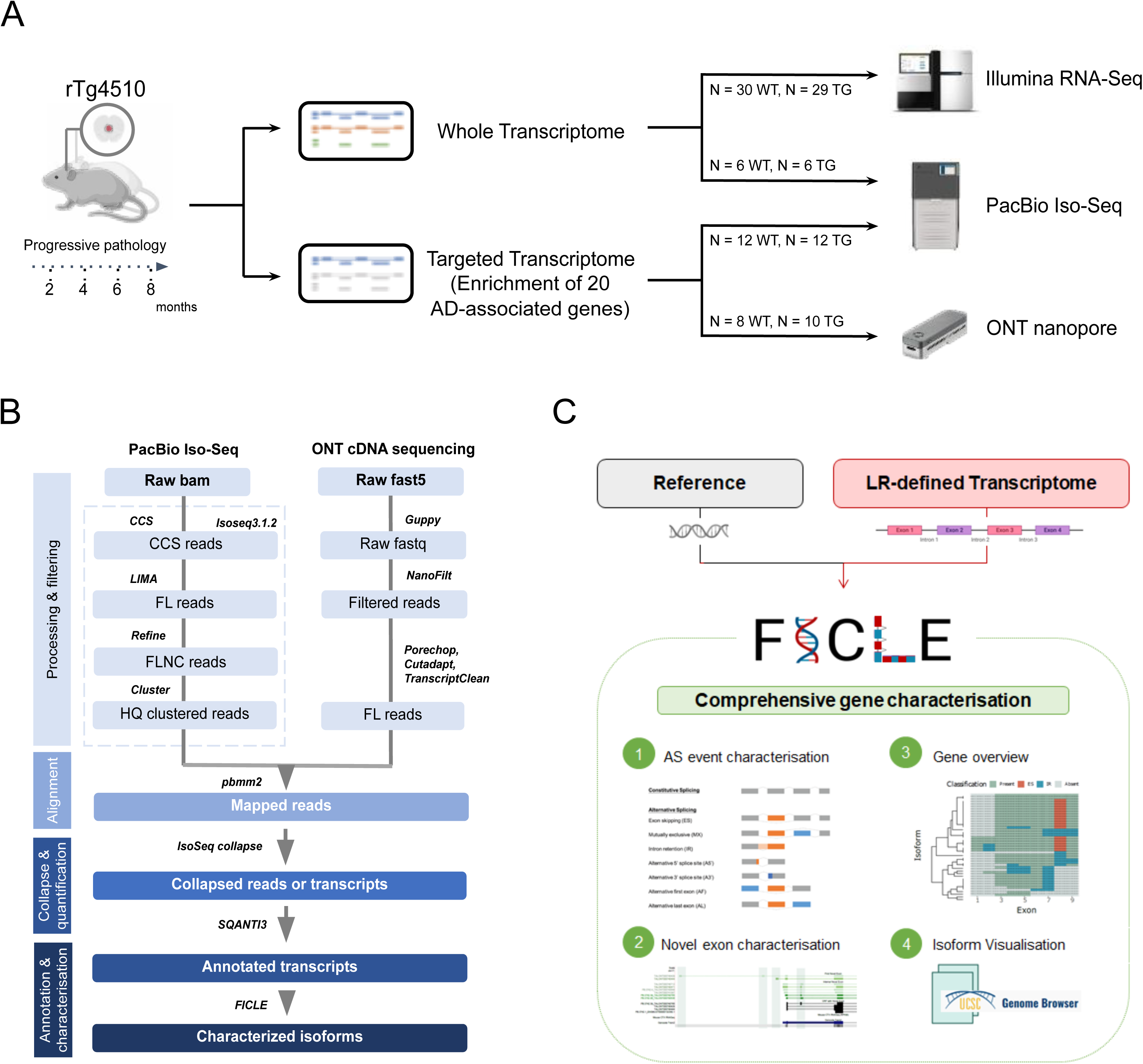
Methodological overview. **(A)** An overview of the experimental approach used to profile isoform diversity in the rTg4510 mouse model using RNA-Seq (Illumina), Iso-Seq (PacBio) and nanopore cDNA sequencing (ONT) at the whole transcriptome and targeted gene level. Further details on the samples can be found in **Methods** and **Supplementary Table 1**. **(B)** An overview of the bioinformatics pipeline used to process, align, quantify, annotate and characterize full-length reads. Further details are provided in **Methods**. **(C)** An overview of the *FICLE* package used to comprehensively characterize and visualize targeted long-read RNA sequencing data.

Across all differentially expressed genes (DEGs) previously associated with the progression of tau pathology in TG mice, we found a strong overall correlation of effect sizes (corr = 0.60, P = 5.17 × 10^−224^) between short-read RNA-Seq and long-read Iso-Seq reads, with a significant enrichment of consistent direction of effects (binomial test: P = 1.45 × 10^−15^, **Supplementary Figure 3**). Our short-read RNA-Seq analysis had previously highlighted *Gfap*, encoding glial fibrillary acidic protein (GFAP), as the top-ranked DEG associated with the progression of tau pathology in TG mice^5^. *Gfap* is highly expressed in astrocytes and up-regulated in reactive astrocytes in response to brain pathology and neurodegenerative disease^11^. Gene-level analysis of our Iso-Seq reads confirmed this association using both normalized full-length Iso-Seq read counts as a proxy for transcript abundance (log_2_ fold change (log2FC) = 3.37, FDR = 1.99 × 10^−4^, **Figure 2A**, **Supplementary Table 3**) and through a ‘hybrid’ approach involving the alignment of short-read RNA-Seq reads from the full set of samples (n = 30 WT, 29 TG at ages 2, 4, 6 and 8 months) to the transcript annotations derived from our Iso-Seq data (log2FC = 2.76, FDR = 2.51 × 10^−17^, **Figure 2B**). We leveraged our full-length Iso-Seq reads to identify a specific differentially expressed transcript (DET) of *Gfap* (LR.Gfap.16, **Figure 2C**) – the most abundantly expressed *Gfap* isoform (Gfap-201, ENSMUST00000067444.9) in the mouse EC – that was the primary driver of overall differential *Gfap* expression in TG mice (log2FC = 3.57, FDR = 1.16 × 10^−3^, **Figure 2D**, **Supplementary Table 4**). The upregulation of LR.Gfap.16 was confirmed using the hybrid method incorporating normalized RNA-Seq read counts (log2FC = 1.06, P = 2.1 × 10^−2^), although this approach also identified upregulation of several novel *Gfap*-associated isoforms (**Figure 2E**). Given the near perfect-homology of these novel isoforms to LR.Gfap.16 (**Figure 2C**), however, we suspect these differences might reflect the limited sensitivity of short RNA-Seq reads for differentiating similar isoforms, highlighting a key advantage of directly using long-read sequencing reads to quantify transcript expression.

**Figure 2:**
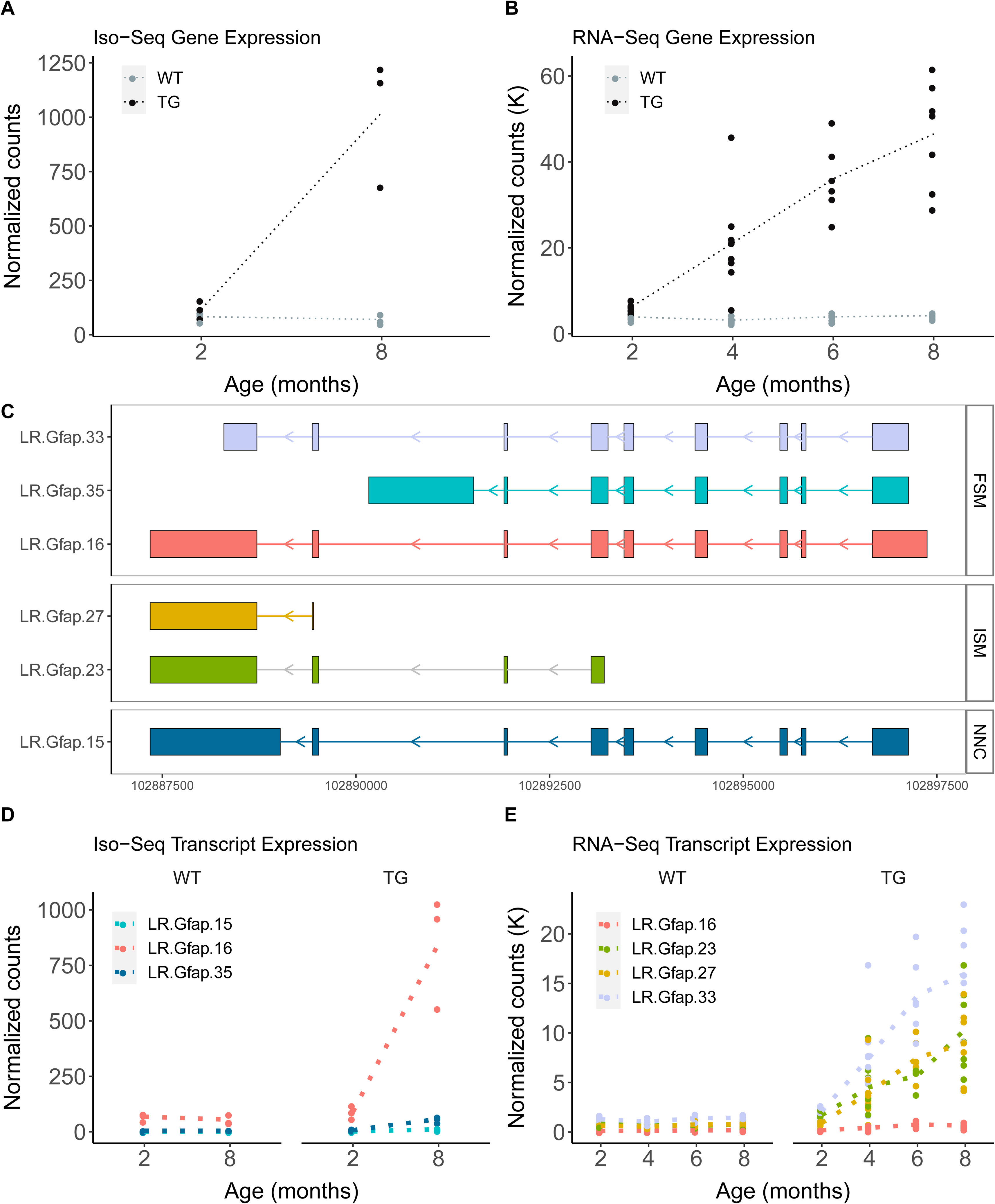
The upregulation of specific isoforms of Gfap is associated with the progression of tau pathology in TG mice. **(A)** *Gfap* gene expression determined from normalized PacBio Iso-Seq full-length read counts and **(B)** normalized short-read RNA-Seq read counts in WT (gray) and TG (black) mice. **(C)** Visualization of the top-ranked differentially expressed *Gfap* transcripts associated with the progression of tau pathology. **(D)** *Gfap* transcript expression determined from full-length Iso-Seq reads and **(E)** RNA-Seq reads aligned to long-read annotations. Colors refer to transcripts in panel **C**. Dotted lines represent the mean paths across ages (months). WT – Wild-type mice, TG – rTg4510 transgenic mice.

### Ultradeep targeted long-read cDNA sequencing identifies hundreds of rare novel isoforms

The constraints on sample numbers (and sequencing depth) possible using the whole transcriptome Iso-Seq approach meant we had relatively limited sensitivity to detect and quantify rare transcripts and identify novel between-group differences using the direct quantification of full-length reads. To enable a comprehensive analysis of alternative splicing and differential transcript usage associated with tau pathology, we therefore performed ultra-deep long-read cDNA sequencing on a targeted panel of 20 genes previously implicated in AD (**Table 1**). We targeted genes affected by autosomal dominant mutations associated with neurodegenerative disorders (*App*, *Mapt*, *Snca*, *Fus*, *Tardbp*), genes identified from both AD genome-wide association studies (GWAS) (*Abca1, Abca7, Apoe, Bin1, Cd33, Clu, Picalm, Ptk2b, Sorl1, and Trem2*) and epigenome-wide association studies (EWAS) (*Ank1, Rhbdf2)*, in addition to several genes implicated in other related neurodegenerative phenotypes (*Fyn, Trpa1, Vgf*). Targeted enrichment was undertaken using custom-designed probes for every known exon across each gene (see **Methods**), followed by sequencing using both ONT nanopore and PacBio Iso-Seq on EC samples from WT and TG mice (see **Figure 1A** and **Supplementary Table 1**).

Our first aim was to characterize the transcriptional landscape for each of the targeted genes in the mouse EC. Raw reads from ONT and Iso-Seq experiments were merged (**Figure 1B**, see **Methods**) and following stringent quality control, our combined dataset comprised of 5.06 million reads annotated to the 20 genes, with no significant difference in read depth between WT and TG mice (mean number of reads per sample: ONT: WT = 248,117 reads, TG = 277,560 reads, Mann-Whitney-Wilcoxon test: W = 50, P = 0.399; Iso-Seq: WT = 12,235 reads, TG = 12,424 reads, Mann-Whitney-Wilcoxon test: W = 81, P = 0.624). As expected, targeted sequencing yielded significantly more unique transcripts of the targeted genes than were detected using whole transcriptome sequencing across the 12 samples processed using both approaches (unique transcripts in targeted = 815, unique transcripts in whole transcriptome = 15, common transcripts in both datasets = 440; fold-change = 2.76, Fisher’s Exact test: P = 4.07 × 10^134^) (**Figure 3A**). We again confirmed expression of human-specific *MAPT* transcripts only from TG mice in the targeted datasets (**Supplementary Figure 2**).

**Figure 3:**
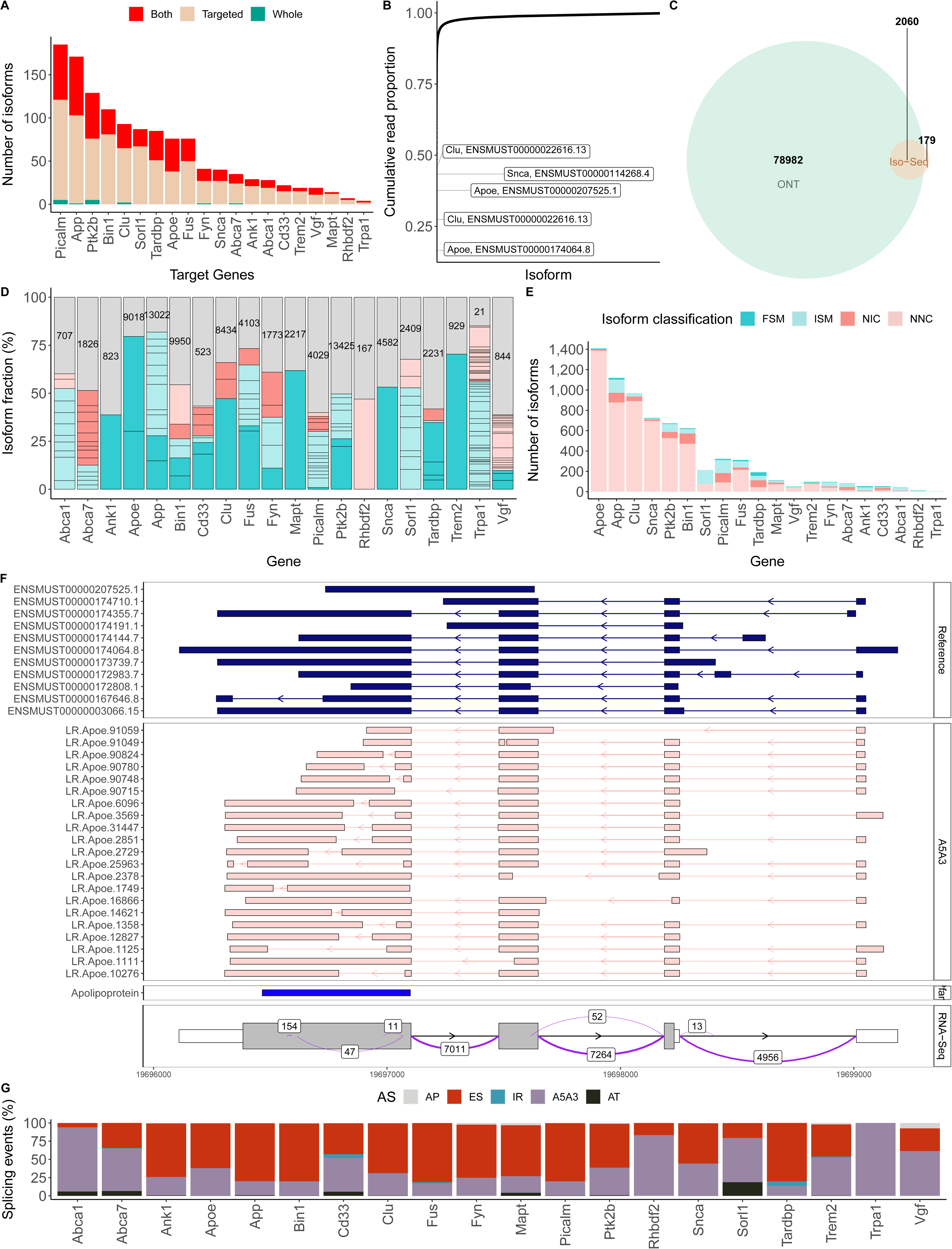
Targeted long-read sequencing identifies considerable isoform diversity and alternative splicing events amongst genes implicated in AD pathology. **(A)** Isoforms detected using whole transcriptome and targeted PacBio Iso-Seq on an overlapping subset of mouse samples (n = 12) across the 20 targeted genes. Commonly detected isoforms using both approaches are depicted in red, and those unique to the targeted and whole transcriptome datasets in beige and green, respectively. **(B)** Cumulative proportion of ONT full-length reads in the targeted dataset. Each dot represents a transcript, and the respective proportion is derived using the summation of reads across all samples (n = 18). **(C)** Venn diagram of the total number of transcripts from the 20 targeted genes detected using targeted ONT (green) and PacBio Iso-Seq (orange) sequencing. **(D)** The isoform landscape and usage across the 20 targeted genes, coloured by *SQANTI3* structural category (FSM - Full Splice Match, ISM - Incomplete Splice Match, NIC - Novel in Catalog, NNC - Novel Not in Catalog). Gray bars refer to “minor” isoforms. The lines across the bars delineate the individual isoforms, and the numbers in the gray bars refer to the total number of minor isoforms. The isoform fraction per gene is determined by dividing the mean normalized read counts of the associated-isoforms across all the samples by the total normalized mean read counts of all the associated isoforms. **(E)** The total number of isoforms annotated to each of the 20 targeted genes, colored by *SQANTI3* structural category (FSM, ISM, NIC, NNC), after merging targeted ONT and PacBio Iso-Seq datasets and filtering by expression (see **Methods**). **(F)** Visualisation of *Apoe* isoforms detected in the merged targeted datasets, highlighting the complexity of the last exon and 3’UTR, as supported by splice junction coverage from short-read RNA-Seq data generated on an extended set of samples (n = 30 WT, 29 TG). **(G)** The proportion of alternative splicing events (AP - alternative promoter, ES - exon skipping, IR - intron retention, A5A3 - alternative 5’ and 3’ splice sites, AT - alternative termination) observed in isoforms annotated to the 20 targeted genes (**Table 1**).

The depth of sequencing achieved using our enrichment approach enabled us to detect 81,221 unique transcripts annotated to the 20 target genes, of which the vast majority were both novel (n = 77,837 (95.8%)) but very rare (74,411 (91.6%) of detected transcripts were characterized by < 10 full-length reads across all samples) (**Supplementary Figure 4**). We detected an average of ~2,200 rare transcripts for each target gene (range = 38 - 13,415) that individually comprised < 1% of sequencing reads, but collectively constituted a relatively large proportion of total reads. Of note, ~50% of the on-target reads were attributed to five transcripts representing known isoforms of *Apoe, Clu* and *Snca* (**Figure 3B**). The majority of novel transcripts were identified in the ultradeep ONT dataset (n = 78,982 transcripts, 97.2%) (**Figure 3C**). Virtually all transcripts identified in the PacBio Iso-Seq dataset (n = 2,060 isoforms, 92%) were also detected by ONT sequencing, whereas a much smaller proportion of transcripts identified in the ONT dataset (n = 2,060 isoforms, 2.54%) were also detected in the PacBio Iso-Seq dataset, reflecting the much higher number of reads obtained from ONT sequencing (**Figure 3C**). Despite the large number of novel transcripts identified, most target genes were characterized by a relatively small number of ‘major’ isoforms (mean = 9.4 isoforms, SD = 10.6, **Figure 3D**). A transcript was defined as ‘major’ if its proportion relative to the dominant (most abundant) isoform was > 0.5 (see **Methods**). This corroborates data from existing datasets such as VastDB – the largest resource of AS events in vertebrates – which shows that the majority of genes were characterized by the simultaneous expression of a relatively small number of major isoforms^12^.

### FICLE - a novel tool for isoform characterization from long-read sequencing data

Existing bioinformatic tools^13,14^ for characterizing AS events using short-read RNA-Seq data fail to capture the connectivity and complexity of transcripts detected from ultra-deep long-read sequencing. Consequently, to comprehensively annotate the full repertoire of transcripts in our data, we developed a novel analysis tool – *FICLE* (see **Methods** and **Figure 1C**) - which we have made available as a resource to the community (see **Code Availability**). *FICLE* directly compares the splice junctions between long-read-derived and reference transcripts, enabling a gene-level overview of all detected isoforms and splicing patterns. It accurately characterizes multiple AS events for each transcript, identifies novel exons, and enables the visualization of isoforms classified by AS events. *FICLE* can be implemented to provide detailed isoform characterization and aid biological interpretations, with the generation of multiple output summary tables and graphs at the exon and transcript level.

### Widespread isoform diversity and alternative splicing events across target genes

After stringent filtering of rare transcripts by removing transcripts with < 10 full-length reads across all samples, our final dataset included 7,162 isoforms comprising 1,000 (14%) known isoforms and 6,162 (86%) novel isoforms. The median number of isoforms across each of the 20 genes was 151, although there was considerable heterogeneity in isoform number between genes. *Apoe* was the most isomorphic gene (1,408 isoforms detected of which 13 were known and 1,395 were novel) whilst *Trpa1* was the least isomorphic gene (4 isoforms detected, all previously characterized) (**Figure 3E**). We used *FICLE* to classify *Apoe* isoforms by exonic structure, observing a complex pattern of AS events involving the last exon, which encodes the apolipoprotein domain that binds to lipids, and the 3’ UTR (**Figure 3F**). Supported by short-read RNA-Seq data from an overlapping set of samples (**Figure 3F**), the variation in the last exon included usage of alternative 5’ and 3’ splice sites and intra-exonic internal skipping, a phenomenon seen in one of the known isoforms (Apoe-202, ENSMUST00000167646.8) (**Figure 3F**). Of note, despite being highly isomorphic, *Apoe* is the shortest gene in our targeted gene panel (gene length = 3.079 kb) with the fewest number of exons (n = 5) (**Supplementary Figure 5**).

For each of the 20 genes, *FICLE* enabled us to identify widespread AS events (**Table 1**, **Figure 3G**). In line with our previous findings^8^, we observed extensive use of alternative 5’ (A5’) and 3’ splice site (A3’) (n = 10,396 events observed in 6,082 (84.9%) isoforms) and exon skipping (ES) (n = 24,315 events observed in 6,031 (84.2%) isoforms) (**Supplementary Figure 6**). Although the majority of isoforms were characterized by the skipping of several exons, some genes expressed transcripts that were characterized by highly elevated levels of ES. For example, over 10% of isoforms expressed from *App* (n = 115 isoforms, 10.2%) and *Bin1* (n = 72 isoforms, 11.5%) were characterized by the skipping of > 10 exons (**Supplementary Figure 7**). In contrast, intron retention (IR) events were less frequent (n = 135 events observed in 134 isoforms, 1.87%) and only detected in isoforms expressed from a subset of the target genes (*Abca7, Apoe, Cd33, Clu, Fus, Ptk2b, Snca, Tardbp* and *Trem2*), corroborating our previous findings^8,15^. Only one isoform (a novel *Cd33* isoform) was characterized with more than one IR event (**Supplementary Figure 7**), although the majority of IR events (n = 126 events, 93.3%) spanned two or more exons with *Tardbp*-associated transcripts demonstrating extensive IR spanning up to 11 exons (**Supplementary Figure 7**).

### Differential transcript expression associated with the progression of tau pathology in TG mice

We focused on our ultradeep ONT sequencing data for the initial identification of specific transcripts associated with the progression of tau pathology in TG mice (see **Methods**), identifying 12 differentially expressed transcripts (DETs) expressed from six of the 20 target genes (*Apoe, App, Cd33, Clu, Fyn*, and *Trem2*) at a FDR < 0.05 (**Figure 4**, **Table 2**, **Supplementary Table 5**). Notably, these transcripts were all up-regulated in TG mice with the progression of tau pathology and five (annotated to *Clu, Fyn*, and *Apoe*) represented novel transcripts not present in existing transcript annotations (**Figure 4**). All 12 of the differentially expressed transcripts were also detected in our PacBio Iso-Seq replication dataset and differentially expressed with the same direction of effect and a strong overall concordance in effect sizes between the two datasets (Log2FC pathology: Spearman’s correlation = 0.643, P = 0.03). Three DETs were confirmed at a Bonferroni-corrected P-value (P < 4.2 × 10^−3^) – a known *Trem2* isoform (LR.Trem2.54), a novel *Clu* (LR.Clu.39341) and novel *App* (LR.App.7564) isoform – and eight additional DETs reached nominal significance (P < 0.05) despite the considerably lower depth of the PacBio sequencing and reduced power for differential expression analyses (**Supplementary Table 5**). We identified an additional 48 DETs expressed from 10 genes (*Abca7, App, Bin1, Cd33, Mapt, Picalm, Ptk2b, Rhbdf2, Snca, Trem2*) associated with genotype (i.e characterized by a significant (FDR < 0.05) difference in expression between WT and TG mice) (**Supplementary Table 6**, **Supplementary Figure 8**). The majority of these DETs (n = 28) were up-regulated in TG mice and a large proportion (n = 14) were annotated to *Trem2* (see below). Of note, the top-ranked up-regulated transcripts in TG mice were mapped to *Mapt (***Supplementary Figure 8**); interestingly, these did not correspond to transcripts derived from the human transgene suggesting the transgene induces concomitant effects on endogenous *Mapt* transcript expression. Among the DETs (n = 31) also detected in the PacBio Iso-Seq dataset, all were again characterized by a consistent direction of effect and there was a strong overall concordance in effect sizes with the ONT data (Log2FC genotype: Spearman’s correlation = 0.792, P = 3.78 × 10^−7^). Finally, we explored differences in *relative* isoform abundance (isoform fraction (IF)) associated with tau pathology, testing for the switching of the dominant (i.e. most highly expressed) isoform between experimental groups using *EdgeR spliceVariants* within *tappAS* (see **Methods**). We identified three genes (*Fus*, *Bin1, Trpa1*) characterized by significant differential transcript usage (DTU) between WT and TG mice, and three genes (*Bin1, Vgf, Clu*) characterized by DTU associated with the progression of tau pathology in TG mice (**Supplementary Figure 9**, **Supplementary Table 7**).

**Figure 4:**
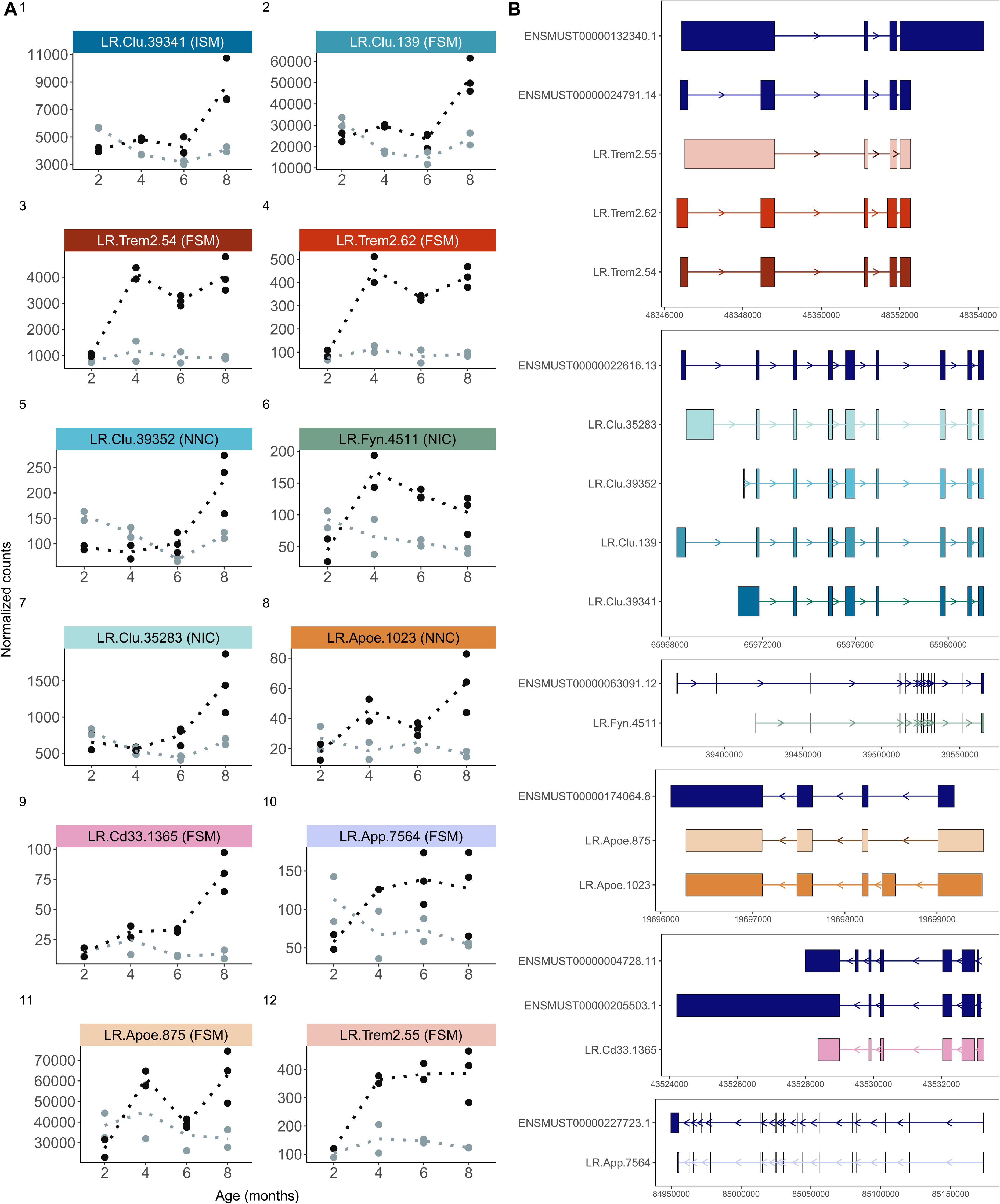
Differential transcript expression associated with the progression of tau pathology in TG mice. **(A)** Transcript expression (ONT FL read counts) of the 12 transcripts significantly up-regulated in TG mice (black) with the progression of tau pathology, and **(B)** visualization of these 12 transcripts with the respective reference transcripts. Plots in panel A are ordered by significance. The colors of the transcripts in panels **A** and **B** are consistent and color-coded by gene, and the shading indicates significance (where darker shades indicate greater significance). *SQANTI3* structural categories (FSM – Full Splice Match, ISM – Incomplete Splice Match, NIC – Novel in Catalog, NNC – Novel Not in Catalog) are provided in parenthesis for each transcript.

**Table 2:**
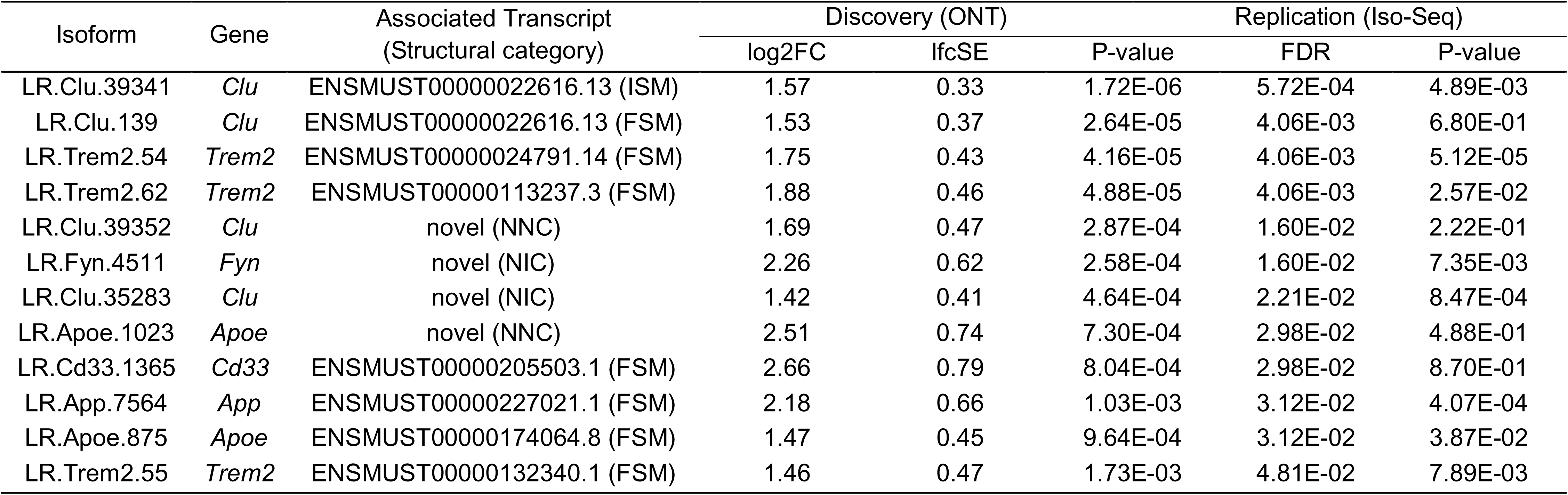
Significant differentially expressed transcripts associated with tau pathology in TG mice. Shown are the 12 differentially expressed transcripts associated with tau pathology in TG mice (**Figure 4**) using the Wald test in *DESeq2* with ONT full-length read counts as a proxy of transcript expression. Results derived from Iso-Seq full-length read counts (targeted PacBio Iso-Seq dataset) are also shown, and the structural categories from *SQANTI3* are also provided. FSM – Full Splice Match, ISM – Incomplete Splice Match, NIC – Novel in Catalog, NNC – Novel Not in Catalog.

### Upregulation of the dominant Trem2 transcript is associated with elevated tau pathology in TG mice

After *Mapt*, the top two DETs in TG mice represented known isoforms of *Trem2* (**Figure 5A**) – LR.Trem2.62 (Trem2-202, ENSMUST00000113237.3) (log2FC = 1.87, FDR = 7.46 × 10^−7^) and LR.Trem2.54 (Trem2-201, ENSMUST00000024791.14) (log2FC = 1.73, FDR = 8.59 × 10^−7^). Both isoforms were also strongly associated with the progression of tau pathology in TG mice (LR.Trem2.62: log2FC = 1.88, FDR = 4.06 × 10^−3^; LR.Trem2.54: log2FC = 1.75, FDR = 4.06 × 10^−3^) (**Figure 4**, **Table 2**). This association was confirmed in our PacBio Iso-Seq targeted dataset with both transcripts associated with genotype (LR.Trem2.62: logFC = 1.73, FDR = 5.2 × 10^−4^; LR.Trem2.54: log2FC = 1.55, FDR = 3.3 × 10^−6^) and pathology in TG mice (LR.Trem2.62: logFC = 2.43, FDR = 9.96 × 10^−1^; LR.Trem2:54: log2FC = 2.20, FDR = 3.85 × 10^−2^). A significant increase of these two known *Trem2* isoforms was also reported in a recent study using qPCR in two transgenic amyloid models of AD^16^,suggesting that these transcripts are upregulated in response to both tau and amyloid pathology. Notably, the up-regulated mouse LR.Trem2.54 isoform shows high homology with the human *TREM2* isoform (ENST00000373113.8, 75.3% homology, 81% query cover) that is most abundantly expressed in the human cortex^17^. Hierarchical clustering of individual samples based on *Trem2* isoform expression level confirmed the robust differences between WT and TG groups across age (**Figure 5B**).

**Figure 5:**
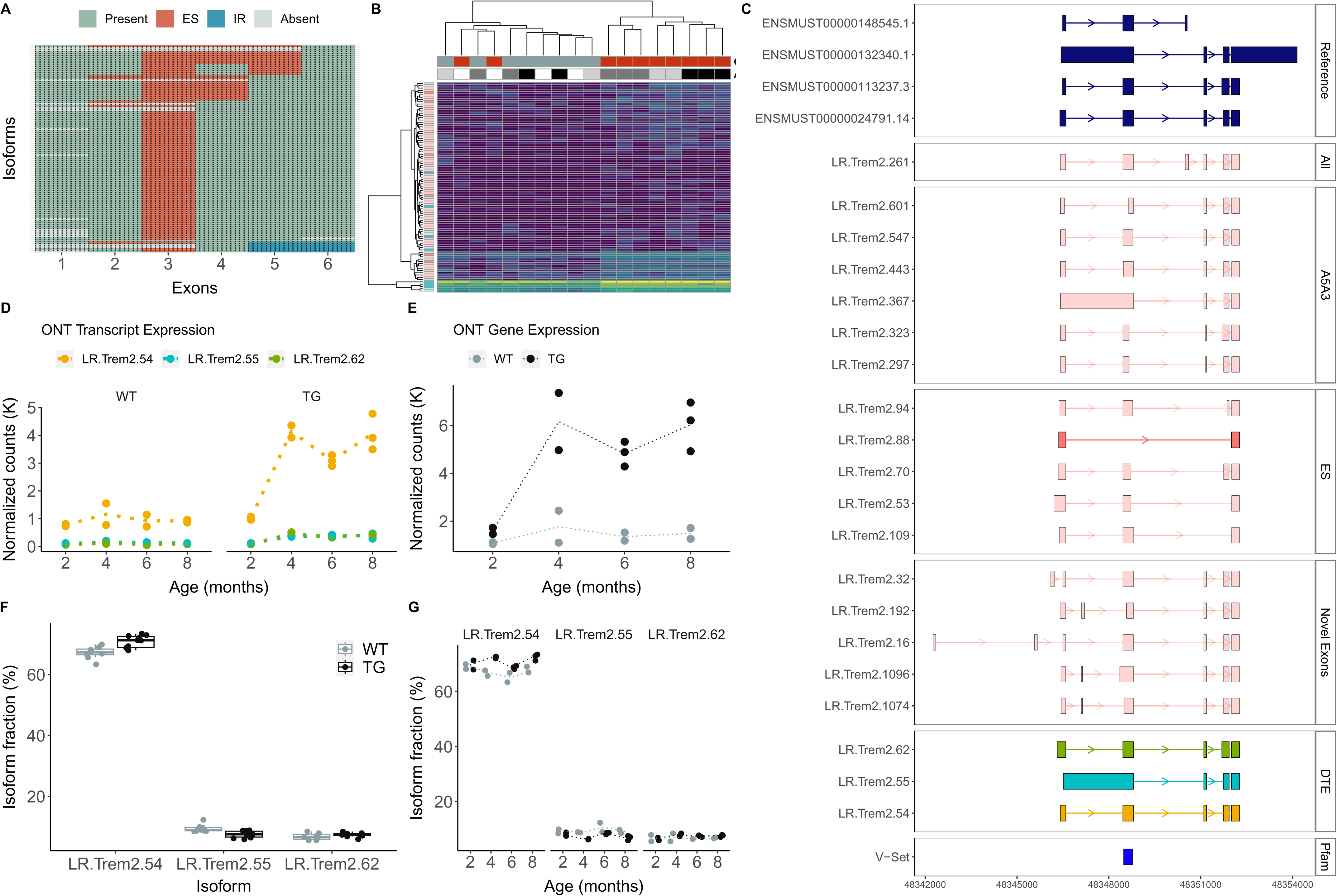
Different isoforms of Trem2 are up-regulated with the progression of tau pathology in TG mice. **(A)** A cluster dendrogram depicting the landscape of *Trem2* isoforms identified using targeted long-read sequencing in the entorhinal cortex from WT and TG mice. Each row corresponds to a distinct isoform and each column represents a known exon. Figure was generated using *FICLE*. **(B)** Heatmap depicting the expression (derived using ONT full-length read counts) of all *Trem2* isoforms after stringent filtering. Each row refers to an isoform, coloured by structural category (FSM – dark blue, ISM – light blue, NIC – pink, NNC – light pink), and each column refers to a sample with the genotype (WT – gray, TG – red) and age (2 months – white, 4 months – light gray, 6 months – dark gray, 8 months – black) provided. **(C)** Visualization of the reference (known) and selected novel *Trem2* isoforms, including a novel isoform incorporating all six known exons (All), novel isoforms with alternative 5’ and 3’ splice sites (A5A3), exon skipping events (ES) and isoforms containing novel exons. Isoforms are coloured by structural category (pink – NIC, light pink – NNC). Shown in the final panel are the three differentially expressed transcripts (DETs) associated with tau pathology, and coloured with reference to panel **D**. **(D)** Transcript expression, determined from normalized ONT full-length read counts, of the three *Trem2* DETs associated with tau pathology (see also Figure 4). **(E)** *Trem2* gene expression (normalized ONT full-length read counts) in WT (gray) and TG (black) mice across age. **(F)** Isoform usage (proportion) of the three *Trem2* DETs by genotype, and **(G)** age.

Overall, *Trem2* is a high isomorphic gene with a large number of novel isoforms (n = 84). However, the expression of the tau-associated DET (LR.Trem2.54) was considerably higher than any of these relatively rare novel isoforms (**Figure 5C, 5D**), suggesting that the upregulation of this isoform is likely to be a primary contributor to the gene-level increase in *Trem2* gene expression observed in older TG mice (log2FC = 2.20, FDR = 3.84 × 10^−2^) (**Figure 5E**). The vast majority of these rare novel isoforms differed primarily in their usage of alternative 5’ and 3’ splice sites (n = 86 isoforms, 88.7%) (**Figure 5A**), particularly in exon 2, as confirmed by short-read RNA-Seq data from matched samples (**Supplementary Figure 10**). This exon was characterized by relatively few ES events (n = 6 isoforms, 6.2%) (**Figure 5A**), potentially reflecting its functional importance; it encodes the Ig-like V-type domain that is crucial for ligand interactions (**Figure 5C**). Of note, the majority of *TREM2* genetic variants associated with AD are located in the homologous region in human *TREM2* isoforms^18^. We also identified 13 *Trem2* isoforms containing novel exons (**Figure 5C**), primarily located between exon 1 and exon 2 with two observed length variants (45bp, chr17: 48347101 - 48347146, n = 5 isoforms; 109bp, chr17: 48347101 - 48347209, n = 2 isoforms). These were retained within the predicted reading frame and may potentially affect protein structure (**Supplementary Figure 10**). Despite the dramatic upregulation of the dominant *Trem2* isoform in TG mice, we observed no overall difference in relative *Trem2* isoform usage between TG and WT mice (**Supplementary Table 7**) with transcript proportions remaining stable between groups (**Figure 5F**) and across age (**Figure 5G**).

### Upregulation of Clu-201 results in differential gene expression and isoform usage

Another gene annotated to multiple DETs associated with the progression of tau pathology in TG mice was *Clu* (**Figure 4**, **Table 2**). The human *CLU* gene encodes a multifunctional glycoprotein that acts as an extracellular chaperone involved in immune regulation and lipid homeostasis^19^. Genetic variants in *CLU* have been associated with late-onset AD^20^. We identified considerable complexity in Clu transcripts with 964 detected isoforms (**Figure 6A, 6B**). In particular there was widespread alternative first exon usage with six distinct alternative first exons, although the majority of *Clu* transcripts (n = 520, 54%) incorporated the first upstream exon of the known dominant isoform (Clu-201, ENSMUST00000022616.13) (**Figure 6A**), skipping these alternative first exons and including the full-length of the clusterin domain (**Figure 6B**). We also found evidence of significant exon skipping and intron retention events localized to other regions of the gene (exon 9 skipping: n = 201 isoforms, exon 10 skipping: n = 225 isoforms, exon 11 skipping: n = 291 isoforms) (**Figure 6A**).

**Figure 6:**
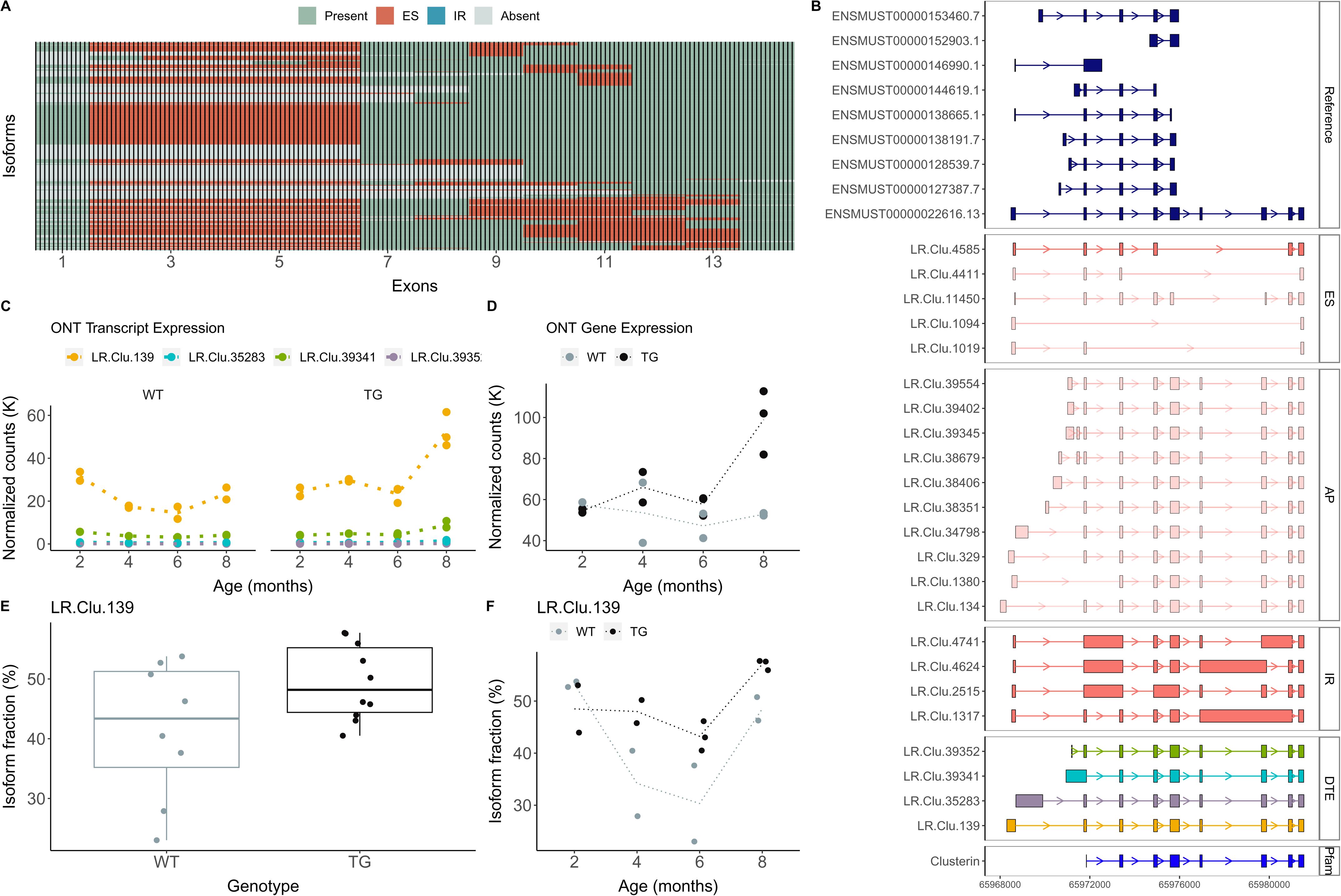
Upregulation of Clu-201 result in differential gene expression and isoform usage. **(A)** A cluster dendrogram depicting the landscape of *Clu* isoforms detected using targeted long-read sequencing in the entorhinal cortex from WT and TG mice. Each row corresponds to a distinct isoform and each column represents a known exon. Figure was generated using *FICLE*. **(B)** Visualization of the reference and selected novel isoforms of *Clu* characterized by exon skipping (ES), intron retention (IR) and alternative promoters (AP). Isoforms are coloured by structural category (pink – NIC, light pink – NNC). Shown in the final panel are the four DETs associated with tau pathology, and coloured with reference to panel **C**. **(C)** Transcript expression, determined from normalized ONT full-length read counts, of the four *Clu* DETs associated with tau pathology (see also Figure 4). **(D)** *Clu* gene expression (normalized ONT full-length read counts) in WT (gray) and TG (black) mice across age. **(E)** Isoform usage (proportion) of LR.Clu.139 by genotype, and **(F)** age.

One of the top-ranked DETs up-regulated with increased tau pathology in TG was LR.Clu.139 (log2FC = 1.13, FDR = 2.73 × 10^−4^) (**Figure 4**, **Figure 6C**), the known dominant isoform Clu-201. Three other *Clu* transcripts that primarily differ from LR.Clu.139 by a novel alternative first exon (**Figure 4B**, **Figure 6B**) were also up-regulated in older TG mice (**Figure 6C**). Upregulation of these four *Clu* transcripts was reflected in an overall increase of *Clu* gene expression observed with increased tau pathology in TG mice (log2FC = 0.97, FDR = 1.1 × 10^−2^) (**Figure 6D**). Of note, there was a notable shift in isoform usage as the proportion of LR.Clu.139 relative to other Clu isoforms (**Supplementary Figure 9**) increased with the progression of pathology in TG mice (total change = 29.2, FDR = 3.12 × 10^−2^) (**Figure 6E, 6F**). These findings corroborate studies on human post-mortem tissue reporting increased expression of the two canonical human *CLU* isoforms in AD, which similarly share a common 3’ exonic structure (exons 2 - 9) but differ in the use of exon 1 and proximal promoters^19^.

### Differential transcript use in Bin1 associated with the progression of tau pathology

Another gene affected by significant DTU was *Bin1*, which was characterized by major changes in isoform proportions associated with both genotype (total change = 17.9, FDR = 2.04 × 10^−2^) and the progression of tau pathology (total change = 30.4, FDR = 7.45 × 10^−14^). The human Bridging Integrator 1 gene (*BIN1*), which is involved in membrane protein trafficking, is another gene strongly implicated in AD genome-wide association studies^20^. We found *Bin1* to be highly isomorphic in the mouse cortex (n = 624 isoforms) with widespread instances of exon skipping that was particularly focused within specific regions of the gene (**Figure 7A, 7B**). As reported for the human *BIN1* gene^21^, the inclusion of exons 14 - 16, which encode the conserved CLAP domain involved in endocytosis^22^, was highly variable among *Bin1* transcripts in the mouse cortex (exon 14 skipping: n = 326 isoforms, exon 15 skipping: n = 242 isoforms, exon 16 skipping: n = 459 isoforms) (**Figure 7B**). In contrast, the first 10 exons of *Bin1*, which encode the N-BAR domain involved in membrane curvature, were relatively conserved and characterized by fewer AS events. Despite the large number of detected *Bin1* isoforms, however, almost a third of ONT *Bin1*-mapped reads were annotated to two isoforms: LR.Bin1.99, the known canonical isoform (Bin1-201, ENSMUST00000025239.8) (n = 16,624 ONT FL reads, 9.4% of *Bin1* ONT FL reads), and LR.Bin1.101, a novel splicing variant of LR.Bin1.99 with a similar exonic structure (n = 36,593 ONT FL reads, 20.7% of *Bin1* ONT FL reads) (**Figure 3D**, **7C**). While the cortical expression of LR.Bin1.99, the canonical isoform, was broadly consistent across genotype groups and age (**Figure 7D**), there was a notable down-regulation at 8 months in TG mice vs WT mice that was paralleled by the significant upregulation LR.Bin1.224, another known isoform (Bin1-205, ENSMUST00000234496.1) (log2FC = 1.19, FDR = 4.78 × 10^−4^) (**Figure 7D**), which differs from LR.Bin1.99 by additional skipping of exon 7 and exon 16 (**Figure 7C**). Notably, we observed differential transcript usage of LR.Bin1.224 between WT and TG mice (WT: mean IF = 4.7%, TG: mean IF = 8.7%, Mann-Whitney-Wilcoxon test: P = 1.18 × 10^−3^) (**Figure 7F**) and with the progression of tau pathology in TG mice (mean IF in TG at 2 months = 6.5%, mean IF in TG at 8 months = 12.7%) (**Figure 7G**), which is likely to be contributing to the overall significant DIU observed in *Bin1* (**Supplementary Figure 9**). These findings corroborate a recent study showing differential *BIN1* isoform expression in the temporal lobe associated with AD in humans^23^. The down-regulated mouse LR.Bin1.99 (Bin1-201) isoform shows high homology with the human *BIN1* isoform 1 (ENST00000316724.10, 87.2% homology, 79% query cover) that is significantly down-regulated in AD, whereas the up-regulated mouse LR.Bin1.224 (Bin1-205) isoform shows high homology to the human *BIN1* isoform 9 (ENST00000409400.1, 88.2% homology, 50% query cover) that is significantly up-regulated in AD (**Supplementary Figure 11**). Despite these striking differences in the abundance of specific *Bin1* isoforms with the progression of tau pathology, there was no overall gene-level expression difference of *Bin1* between WT and TG mice (log2FC = −0.079, P = 0.65) (**Figure 7E**), further highlighting the importance of performing transcript-level analyses.

**Figure 7:**
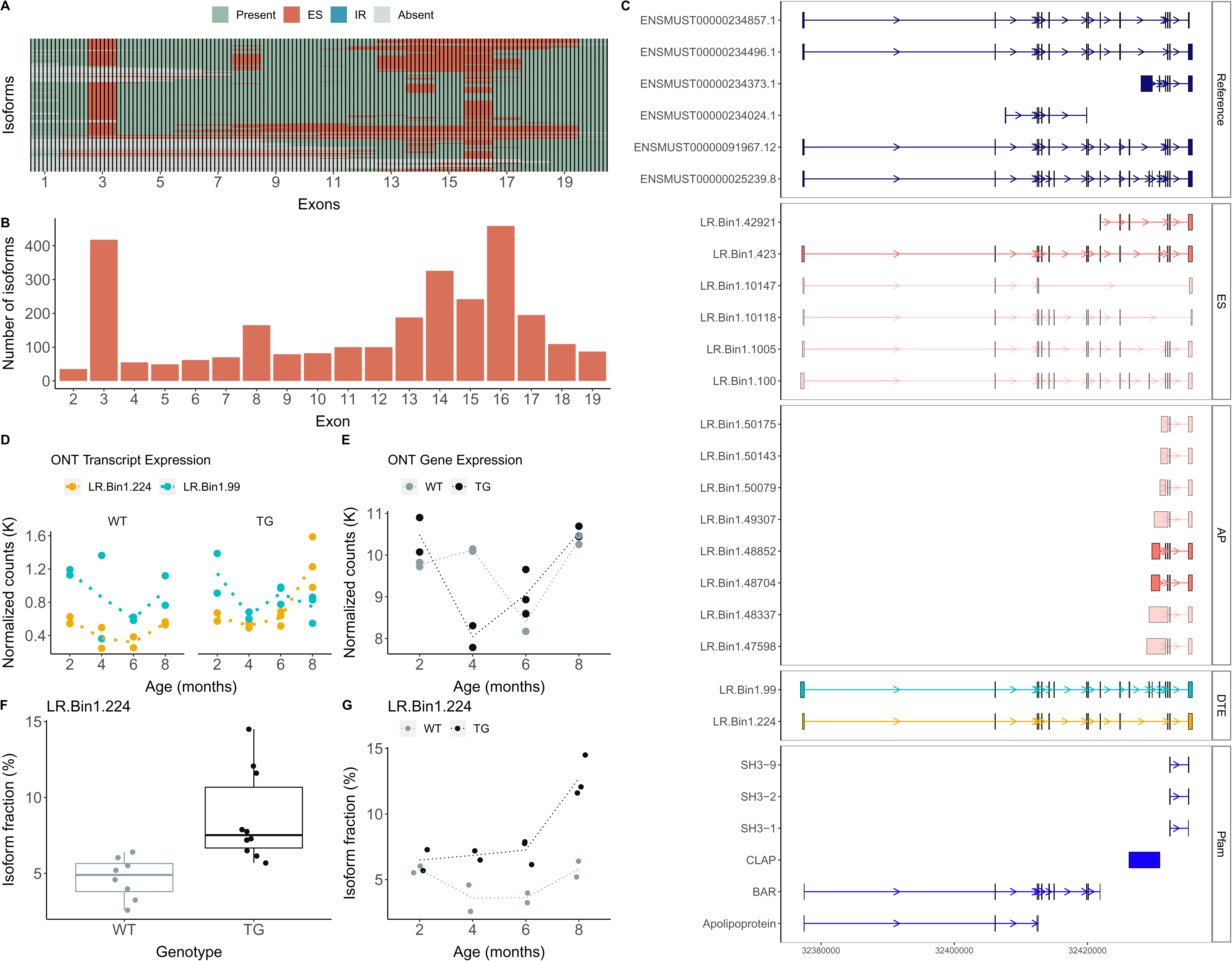
Isoform switching in *Bin1* associated with tau pathology in TG mice. **(A)** A cluster dendrogram depicting the landscape of *Bin1* isoforms detected using targeted long-read sequencing in the entorhinal cortex from WT and TG mice. Each row corresponds to a distinct isoform and each column represents a known exon. **(B)** A box-plot of the number of isoforms in which the known exon (mm10, GENCODE) is skipped. The numbering of the known exon is determined from flattening the reference transcripts (mm10) using *FICLE*. **(C)** Visualization of the reference and selected novel isoforms of *Bin1* characterized by exon skipping events (ES), intron retention (IR) and alternative promoter use (AP). Isoforms are coloured by structural category (pink – NIC, light pink – NNC). **(D)** Transcript expression, determined from normalized ONT full-length read counts, of two known isoforms: LR.Bin1.99, (Bin1-201, ENSMUST00000025239.8) and LR.Bin1.224 (Bin1-205, ENSMUST00000234496.1). Isoforms are coloured with reference to panel **C**. **(E)** *Bin1* gene expression (normalized ONT full-length read counts) in WT (gray) and TG (black) mice across age. **(F)** Isoform usage of LR.Bin1.224 by genotype, and **(G)** age.

## Discussion

We leveraged the power of long-read sequencing to characterize transcript diversity in the mouse cortex and identify differentially expressed transcripts (DETs) associated with the progression of tau pathology. Our study represents the first analysis of isoform diversity using long-read cDNA sequencing in a transgenic model of AD pathology and highlights the importance of differential transcript usage, even in the absence of gene-level expression alterations, as a potential mechanism underpinning the consequences of pathogenic tau deposition. Ultra-deep targeted long-read sequencing of genes previously implicated in AD revealed hundreds of novel isoforms driven by AS events and transcript level splicing differences paralleling the development of tau pathology and reflecting changes observed in human AD brain tissue. Our transcript annotations and novel tool for the characterization of long-read transcript sequencing data (*FICLE*) are provided as a resource to the research community.

Our whole transcriptome analysis confirmed findings from our previous analysis of rTg4510 mice using short-read RNA-Seq^5^ revealing highly-similar gene-level expression differences associated with tau pathology in TG mice. We were able to leverage our full-length transcript reads to identify the specific transcripts driving these gene-level differences; for example, we confirmed the dramatic upregulation of *Gfap* and showed that this was primarily driven by the increased abundance of the canonical Gfap-201 isoform.

To enable a more comprehensive analysis of alternative splicing and differential transcript use associated with tau pathology, we performed ultra-deep long-read cDNA sequencing on a targeted panel of 20 genes previously implicated in AD using custom-designed biotinylated probes to enrich full-length transcripts prior to both ONT nanopore sequencing and PacBio Iso-Seq. We identified thousands of alternatively-spliced isoforms and detected many novel transcripts annotated to these genes highlighting the power of targeted cDNA sequencing. We established *FICLE* – a tool to handle and document the complexity of these isoforms detected from high-throughput long-read sequencing – which we subsequently used to comprehensively characterize AS events.

Our findings highlight the extent to which AS events contribute to isoform diversity in the cortex and highlight the power of long-read sequencing approaches for transcriptional profiling. The deep sequencing depth achieved using target enrichment allowed us to reliably identify DETs using normalized full-length read counts derived from long-read sequencing. We identified widespread transcriptional variation associated with the progression of tau pathology in TG mice with evidence of altered splicing and transcript expression across the 20 AD genes targeted in our experiments. We identified tau-associated DETs annotated to six of the 20 target genes (*Apoe, App, Cd33, Clu, Fyn*, and *Trem2*). We also identified evidence of DTU associated with tau pathology in genes not characterized by an overall difference in gene expression, highlighting the power of long-read sequencing for characterizing transcriptional differences not detectable using traditional short-read RNA-Seq approaches. Many of the differences observed parallel changes reported in studies of AD in the human cortex. For example, we find evidence of differential *Bin1* isoform use in older TG mice paralleling findings from a study in human AD post-mortem cortex^23^. We show that the differentially expressed isoforms primarily differed by the presence/absence of exons encoding the clathrin-binding domain (CLAP) domain, which is involved in endocytosis and is also highly variable among human-equivalent *BIN1* isoforms^21^. Our results further implicate altered exon splicing as a potential mechanism contributing to the role of *Bin1* in tau pathology.

Our results should be interpreted in the context of several limitations. First, because the PacBio Iso-Seq protocol does not include 5’-cap selection we cannot exclude the possibility that some of the shorter isoforms identified in our study result from 5’ degradation. By using two long-read sequencing platforms (ONT and PacBio) to characterize isoforms in the mouse cortex, in addition to short-read RNA-seq, we were able to validate a large proportion of our isoform annotations and reduce the number of these artifacts. Of note, many of the novel transcripts identified in this study were rare and excluded from downstream association analyses. Second, our analyses were performed on ‘bulk’ entorhinal cortex tissue, comprising a heterogeneous mix of neurons, oligodendrocytes and other glial cell-types. Despite compelling evidence from recent studies reporting cell-specific transcriptional signatures in disease^24^, we were unable to explore these differences in this study. Of note, novel approaches for using long-read sequencing approaches in single cells will enable a more granular approach to exploring transcript diversity in the cortex^25^. Furthermore, while isoform expression was normalized for potential technical cofounders (e.g. library sequencing depth), it was not possible to account for differences in cellular composition between WT and TG mice. Given neuronal loss and astrogliosis are prominent hallmarks of AD pathogenesis, we were unable to discern whether transcript-level variations are a direct consequence of AD-associated transcriptional regulation or reflect changes in cell composition associated with pathology. Finally, we only profiled entorhinal cortex tissue given that this is one of the first regions of the brain to be affected by AD pathology However, tissue-specific differences in splicing and isoform usage of AD-risk genes have been previously reported^26^. Furthermore, in order to reduce heterogeneity in our analyses we only profiled tissue from female mice; a number of sex differences have been reported, with female mice exhibiting earlier and more severe cognitive and behavioral impairments than male TG mice^27^. Future work would cross-examine results from our study with transcriptional variation in other tissue types and male mice for a more comprehensive understanding of the development of tau pathology in the AD brain.

In summary, our study identified transcript-level differences in the entorhinal cortex associated with the accumulation of tau pathology in TG mice. Importantly, we identified changes in the abundance of specific transcripts that drive the altered expression of AD genes, with evidence of isoform switching events that could have important functional consequences. Taken together, our results demonstrate the utility of long-read sequencing for transcript-level analyses, facilitating the detection of AS events associated with the development of AD pathology. We have made our transcript annotations and analysis pipeline freely available as a resource to the community to stimulate further research in this area.

## Acknowledgements

S.K.L. was supported by a UK Medical Research Council (MRC) CASE PhD studentship. This work was funded in part through the MRC Proximity to Discovery: Industry Engagement Fund (Precision Medicine Exeter Innovation Platform reference MC_PC_14127) and a research grant from Alzheimer’s Research UK (ARUK-PG2018B-016). I.C.’s doctoral studentship was supported by the Alzheimer’s Society in partnership with the Garfield Weston Foundation (grant 231). Development of *FICLE* was supported by a grant from the Simons Foundation for Autism Research (SFARI) (grant number 573312, awarded to J.M.). Sequencing and computational facilities were supported by an MRC Clinical Infrastructure award (MR/M008924/1); the Wellcome Trust Institutional Strategic Support Fund (WT097835MF); a Wellcome Trust Multi-User Equipment Award (WT101650MA); a Biotechnology and Biological Sciences Research Council (BBSRC) Longer and Larger (LoLa) award (BB/K003240/1) and the University of Exeter High-Performance Computing (HPC).

## Author contributions

S.K.L. and A.R.J. conducted long-read sequencing experiments. A.R.J., I.C. and K.M. conducted short-read RNA-Seq experiments. K.M. advised on library preparation and aspects of sequencing. Z.A. provided mouse cortex tissue. J.M., E.H. and E.L.D. obtained funding. J.M. and S.K.L. designed the study. S.K.L. undertook primary data analyses and bioinformatics, with analytical and computational input from A.R.J, P.O. and E.H. R.B., Z.A., J.T.B and E.L.D helped interpret the results. S.K.L. and J.M. drafted the manuscript. All authors read and approved the final submission.

## Methods

### Samples

Mouse entorhinal cortex tissue was dissected from female wild-type and rTg4510 transgenic (TG) age-matched littermate mice in accordance with the UK Animals (Scientific Procedures) Act 1986 and with approval of the local Animal Welfare and Ethical Review Board. Details on breeding conditions and sample preparation can be found in Castanho *et al.* 2020^5^. Entorhinal cortex tissue was dissected from 29 female rTg4510 TG and 30 female WT mice, aged 2, 4, 6 and 8 months (n = 7 - 8 mice per group).

### RNA isolation and highly-parallel RNA sequencing

Short-read RNA-Seq data was previously generated for each mouse to identify differentially expressed genes associated with the progression of tau pathology^5^. Briefly, RNA was isolated using the AllPrep DNA/RNA Mini Kit (Qiagen, UK) from ~5 mg tissue and quantified using the Bioanalyzer 2100 (Agilent, UK). RNA-Seq libraries were subsequently prepared with the TruSeq Stranded mRNA Sample Prep Kit (Illumina) and subjected to 125 bp paired-end sequencing using a HiSeq2500 (Illumina). Raw sequencing reads were then processed and mapped to the mouse (mm10) reference genome using *STAR*^44^ (v1.9). RNA-Seq-derived gene and transcript expression were then determined by aligning RNA-Seq reads to the whole transcriptome Iso-Seq dataset (*SQANTI3* filtered) generated in this study using *Kallisto*^45^ (v0.46.0).

### Whole transcriptome Iso-Seq library preparation and SMRT sequencing

Whole transcriptome PacBio Iso-Seq was performed on RNA isolated from a subset of 12 mice (3 WT and 3 TG at ages 2 and 8 months) (**Supplementary Table 1**). First strand cDNA synthesis was performed on 200ng RNA using the SMARTer PCR cDNA Synthesis Kit (Clontech, UK) according to manufacturer’s instructions. A total of 14 PCR cycles of amplification was performed for each sample using PrimeSTAR GXL DNA Polymerase (Clontech, UK). Library preparation of the amplified cDNA products was then performed using SMRTbell Template Prep Kit v1.0 (PacBio, USA). Sequencing was performed on the PacBio Sequel 1M SMRT cell. Samples were processed using either the version 3 chemistry (diffusion loading at 5 pM, 4 hours pre-extension, 20 hours capture) or version 2.1 chemistry (magbead loading at 50 pM, 2 hours pre-extension, 10 hours capture).

Raw reads were processed and de-multiplexed with *Iso-Seq* (v3), aligned to the mouse reference genome (mm10, GENCODE) using *pbmm2*^46^ (v1.10.0, a *Minimap2* wrapper adapted for PacBio data, parameters: --preset ISOSEQ --sort), collapsed to full-length transcripts using *Iso-Seq collapse*^47^ (v3.8.2) and annotated using *SQANTI3* (v5.0) in combination with the mouse reference gene annotations (mm10, GENCODE, vM22). SQANTI3 filtering was performed using the default JSON file, with the sole modification of relaxing the “min_cov” (the minimum short-read RNA-Seq coverage threshold of non-canonical junctions) from 3 to 0 for non-FSM isoforms. Given the extensive coverage achieved by our targeted long-read sequencing data and the comparably lower depth of our short-read RNA-Seq data for the target genes, we concluded that this parameter was redundant. Full-length Iso-Seq read counts from each sample were extracted from the *Iso-Seq collapse* “read_stat.txt” file with the CCS read ID as sample identifiers.

### Targeted Iso-Seq library Preparation and SMRT sequencing

Targeted PacBio Iso-Seq was performed on RNA isolated from a subset of 24 mice (3 WT and 3 TG at ages 2, 4, 6 and 8 months) (**Supplementary Table 1**). The same WT and TG mice at aged 2 and 8 months were previously sequenced using whole transcriptome PacBio Iso-Seq (**Supplementary Table 1**). First-strand cDNA synthesis was performed on 200ng RNA using the SMARTer PCR cDNA Synthesis Kit (Clontech) with specific oligo(dT) barcodes for multiplexing. Large-scale PCR amplification was subsequently performed using 14 cycles, and the resulting amplicons were subjected to targeted enrichment using custom-designed probes (IDT, UK). To avoid off-target binding, we manually assessed the list of probes for each target gene under the following criteria: i) all of the exons must be covered by at least one probe, ii) probes should be spaced 300bp - 500bp within each exon (equivalent to 0.2x – 0.3x tiling density), iii) probes with the highest GC content (40 - 65% GC content) and lowest number of blast hits were selected from the contiguous cluster, and iv) any probes covering the intronic regions were removed. Following successful enrichment for target genes, Iso-Seq library preparation was performed using the SMRTbell Template Prep Kit v1.0 (PacBio) for subsequent sequencing on 3 PacBio Sequel 1M SMRT cells. Raw reads were processed and de-multiplexed with *Iso-Seq* (v3).

### Targeted ONT library preparation and nanopore sequencing

Ultradeep targeted ONT nanopore sequencing was performed on RNA isolated from a subset of 18 mice (8 WT and 10 TG at ages 2, 4, 6 and 8 months). All mice were previously sequenced using targeted PacBio Iso-Seq (**Supplementary Table 1**). ONT library preparation was undertaken using the Ligation Sequencing Kit (SQK-LSK109, ONT) after target enrichment as described above. Sequencing was subsequently performed on the ONT MinION using two FLO-Min106D flow cells. Raw ONT reads were then basecalled using *Guppy* (v4.0) and reads with Phred (Q) < 7 were discarded. Primers and ONT adapters were removed using *Porechop* (v0.2.4) to generate full-length reads for each sample. After trimming of poly(A) tails using *Cutadapt* (v2.9), full-length reads were then aligned to the mouse reference genome (mm10, GENCODE) using *Minimap2* (v2.17, parameters: “-ax splice”) and corrected using *TranscriptClean*.

### Merged annotation and quantification from targeted long-read sequencing datasets

To comprehensively characterize the transcripts annotated to each of the 20 target genes, full-length reads from both targeted Iso-Seq (after *Iso-Seq3* cluster) and ONT datasets (after correction with *TranscriptClean*) were merged. The merged dataset was then aligned to the mouse reference genome (mm10, GENCODE, vM22) using *pbmm2*^46^ (v1.10.0) and collapsed to full-length transcripts using *Iso-Seq collapse*^47^ (v3.8.2). Full-length Iso-Seq and ONT read counts from each sample were extracted using custom scripts that determined the source of the reads clustered from the *Iso-Seq collapse* “read_stat.txt” output file. The merged dataset was then annotated with *SQANTI3* in combination with the mouse reference gene annotations (mm10, GENCODE, vm22). *SQANTI3* filtering was performed using the default JSON file, with the sole modification being the decrease of “min_cov” from 3 to 0 for non-FSM isoforms. Isoforms associated with the 20 target genes were subsequently selected and retained if observed more than 10 times (i.e. full-length read count ≧ 10) across any five samples (**Supplementary Figure 4**).

### Characterisation of AS events and transcript visualization

We developed a novel python-based tool, *FICLE* (v1.1.2), to accurately assess the occurrence of alternative splicing events by comparing splice sites (exon) coordinates between long-read-derived transcripts and reference transcripts (GENCODE). Common alternative splicing events including alternative first exon use (AF), alternative last exon use (AL), alternative 5’ splice sites (A5), alternative 3’ splice sites (A3), intron retention (IR) and exon skipping (ES) were assessed. Alternative 5’ and 3’ splice sites were defined as splice sites differing by more than 10bp from the known splice site. Other regulatory mechanisms such as alternative transcription initiation (defined by an alternative transcription start site) with the presence of an alternative promoter (AP) and termination (defined by an alternative transcription termination site) with the presence of an alternative terminator (AT), and the presence of novel exons not present in existing transcript annotations, were also evaluated. Open reading frames were predicted using the Coding-Potential Assessment Tool^48^ (CPAT) (v3.0.2) with default parameters, and transcripts with coding potential score > 0.44 (recommended threshold for mouse) were predicted as protein-coding. Isoforms were visualized using either the UCSC genome browser or *ggtranscrip*t (v0.99.9)^49^.

### Differential expression and splicing analyses

Differential expression analysis was performed using *DESeq2*^50^ (v1.26.0) with Iso-Seq and ONT full-length read counts as proxies of gene and transcript expression. Notably, gene expression was aggregated as the summation of full-length read counts from all isoforms associated with target genes, prior to filtering for rare (lowly-expressed) transcripts (minimum 10 reads across any 5 samples). Briefly, DESeq2 normalizes read counts using the median of ratios method, estimates dispersion, and tests for differential expression using negative binomial generalized linear models. Datasets were filtered for lowly expressed isoforms (minimum of 10 reads across all samples). Significant genotype effects between WT and TG mice were identified using the Wald test: ~ genotype. Significant interaction effects to detect progressive changes across age between WT and TG mice (i.e. pathology) were identified using the Wald test to compare the nested regression model: ~ genotype + age + genotype * age. Significance was defined by FDR < 0.05, after adjusting *P values* for multiple testing using the false discovery rate (FDR) method (also known as Benjamini and Hochberg correction).

Differential transcript usage analysis was performed using *EdgeR spliceVariant*^51^ (v3.28.1) within *tappAS*^52^ (v1.0.7). Lowly-expressed isoforms were filtered using *minorFoldfilterTappas*, whereby an isoform was only retained if its relative proportion to the major isoform was above a specified fold change (FC) threshold (default FC = 0.5). Adopting *tappAS* DIU analysis and metrics, we were able to quantify the magnitude of redistribution of gene expression across its isoforms between conditions (total change) and detect major isoform switching events (podium change). To illustrate DIU at an isoform and sample level (for example, **Figure 5F** and **Figure 5G**), the isoform usage was determined by dividing the normalized read count for each associated isoform by the total normalized read counts of all the associated isoforms. To illustrate DIU at an isoform level only (**Figure 3D**), the isoform usage was determined by dividing the mean normalized read count for each associated isoform across all the samples by the total normalized read counts of all the associated isoforms.

## Data and Code Availability

Raw ONT and PacBio Iso-Seq data has been deposited in the Sequence Read Archive (SRA) database (https://www.ncbi.nlm.nih.gov/sra) under accession numbers PRJNA981131 (targeted ONT and PacBio Iso-Seq data) and PRJNA663877 (whole transcriptome Iso-Seq data). Intermediate files and UCSC genome browser tracks (merged targeted data and whole transcriptome Iso-Seq data) are available at: https://doi.org/10.5281/zenodo.8101908. All original code supporting this study is available at https://github.com/SziKayLeung/rTg4510, and the *FICLE* package is available to download at https://github.com/SziKayLeung/FICLE.

