## Supplementary Figure 1-11 for "Long-read transcript sequencing identifies differential isoform expression in the entorhinal cortex in a transgenic model of tau pathology"

**Supplementary Figures**

| Supplementary Figure 1 | Global transcript characteristics between WT and TG mice |
| --- | --- |
| Supplementary Figure 2 | Presence of human MAPT transgene in TG mice |
| Supplementary Figure 3  Supplementary Figure 4  Supplementary Figure 5 | Correlation of RNA-Seq and Iso-Seq gene expression  Rarefaction curve to determine expression threshold  Relationship between number of isoforms and other features |
| Supplementary Figure 6 | Characterisation of ES and A5’A3’ events in AD-associated genes |
| Supplementary Figure 7  Supplementary Figure 8  Supplementary Figure 9  Supplementary Figure 10  Supplementary Figure 11 | Characterisation of IR in AD-associated genes  Top 10 differentially expressed transcripts (genotype)  Differential transcript usage  Further characterization of *Trem2*  Homology of *Bin1/BIN1* isoforms in mouse and human |

**Supplementary Figure 1: Global transcript characteristics between WT and TG mice**.

Shown are box-plots of the **(A)** isoform length and the **(B)** exon diversity of the isoforms commonly (“Both”) and uniquely detected in WT and TG mice from the whole transcriptome PacBio Iso-Seq dataset. There was no difference between groups for any measure.

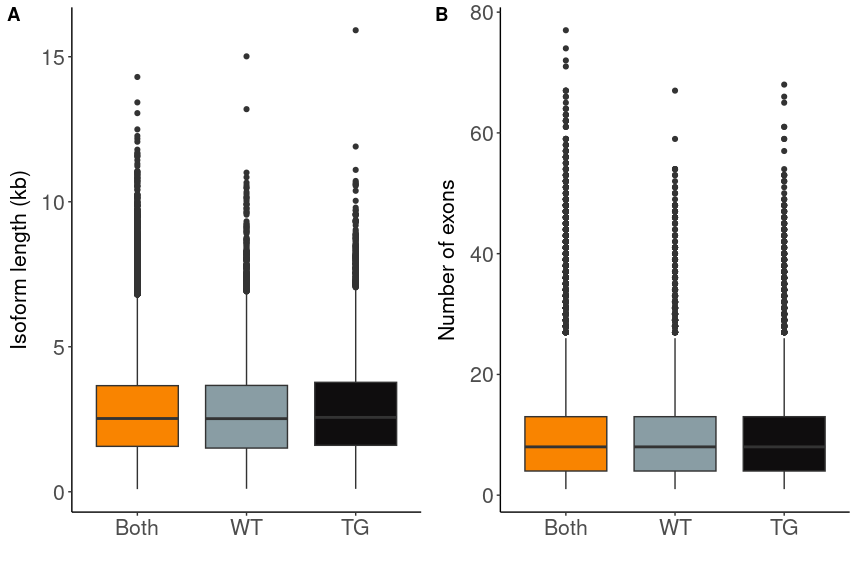

**Supplementary Figure 2: Human-specific *MAPT* sequences were only present in transgenic mice**.

Shown are scatter plots of the proportion of full-length transcripts that were mapped to human-specific *MAPT* and mouse-specific *Mapt* sequences in the **(A)** whole transcriptome PacBio Iso-Seq, **(B)** targeted PacBio Iso-Seq, and **(C)** targeted ONT datasets. Red and gray dots refer to TG and WT samples, and dotted lines represent the mean paths across ages.

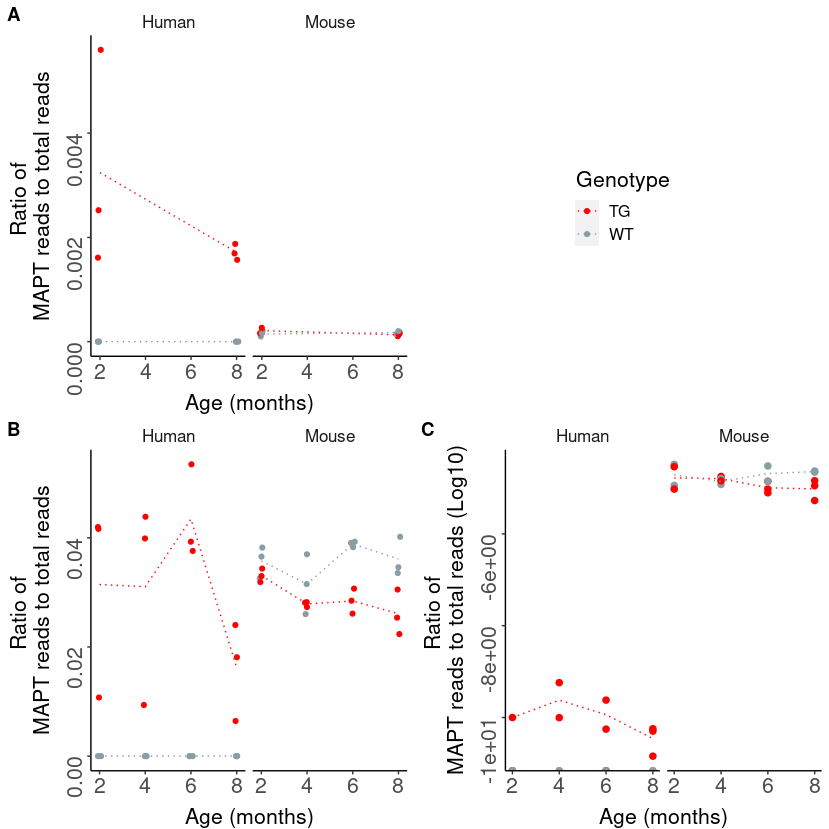

**Supplementary Figure 3: Concordant gene expression differences observed between whole transcriptome Iso-Seq and RNA-Seq datasets.**

Shown is a strong positive correlation of DEG effect sizes (log_2_ fold change between WT and TG at 8 months) derived from Iso-Seq full-length reads (whole transcriptome Iso-Seq dataset) and short-read RNA-Seq reads (RNA-Seq dataset) as a proxy of gene expression in *DESeq2*. Each dot represents a gene that is differentially expressed between WT and TG mice with the progression of tau pathology (pathology effect) in the RNA-Seq (n = 59) dataset (Castanho et al. 2020), following alignment to transcript annotations derived from our Iso-Seq data.

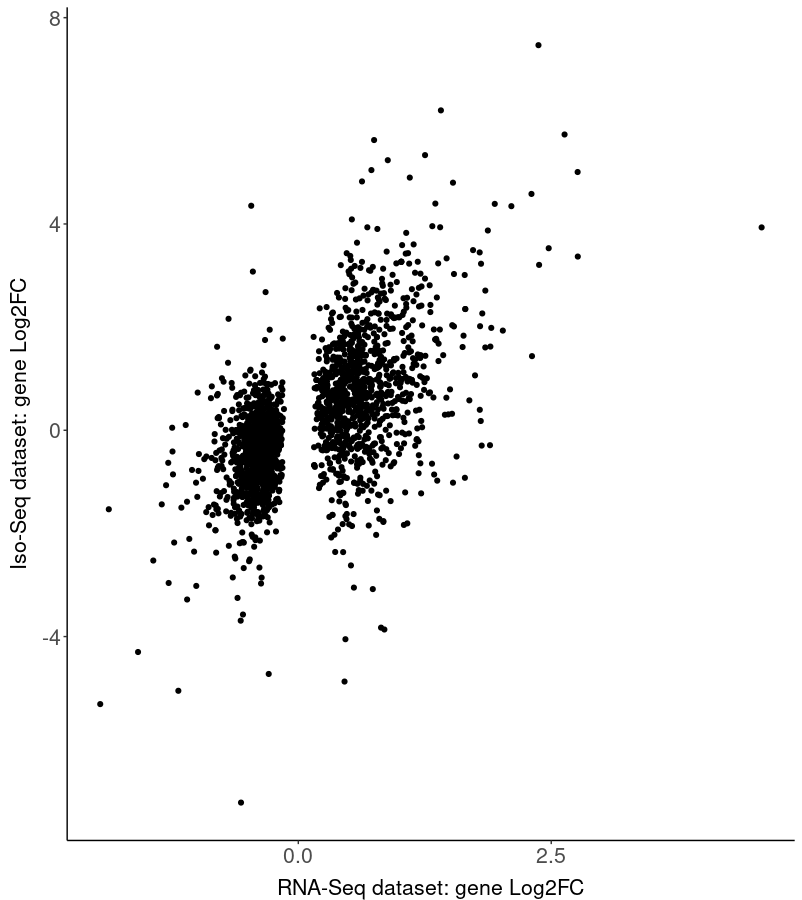

**Supplementary Figure 4: Rarefaction curve to determine expression threshold for filtering targeted datasets.**

Shown is a rarefaction curve of the percentage of transcripts retained after applying stepwise sample and read-count thresholds. For example, the red data points on the left-to-right X-axis represent the percentage of transcripts (associated to the 20 target genes from merged targeted ONT and Iso-Seq datasets) retained after applying thresholds of (total) full-length reads ≥ 2, 3, …10, 15, and 20 reads across any 2 samples. By identifying the point at which all the curves start to converge, we established a minimum expression threshold of 10 reads across at least 5 samples for filtering rare transcripts in our targeted datasets.

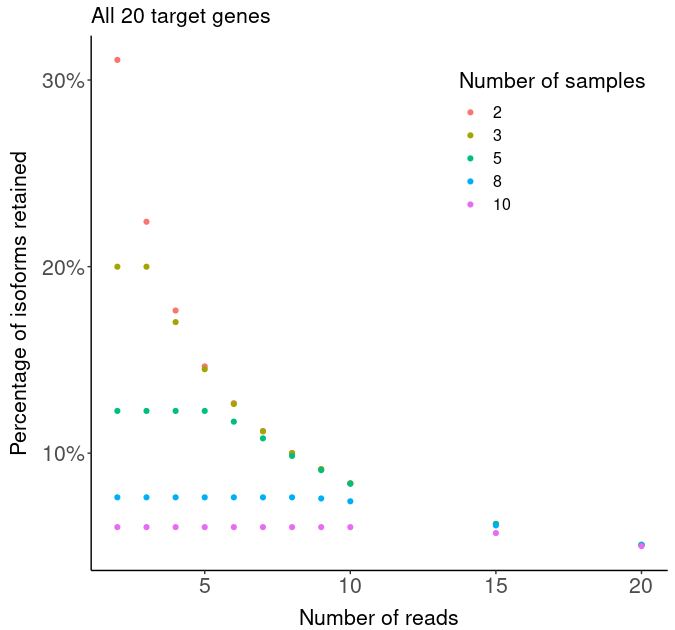

**Supplementary Figure 5: Relationship between the number of isoforms and other features in targeted ONT dataset.**

Shown are scatter plots of the number of detected isoforms against **(A)** gene length, **(B)** the number of exons, and **(C)** the normalized count as a proxy of gene expression. Each dot refers to a target gene from the targeted ONT dataset. The gene length and the number of exons (maximum number) are extracted from reference mouse annotations (mm10, GENCODE). **(D)** Distribution of isoforms detected by abundance across the target genes from the targeted ONT dataset. Each row refers to an isoform.

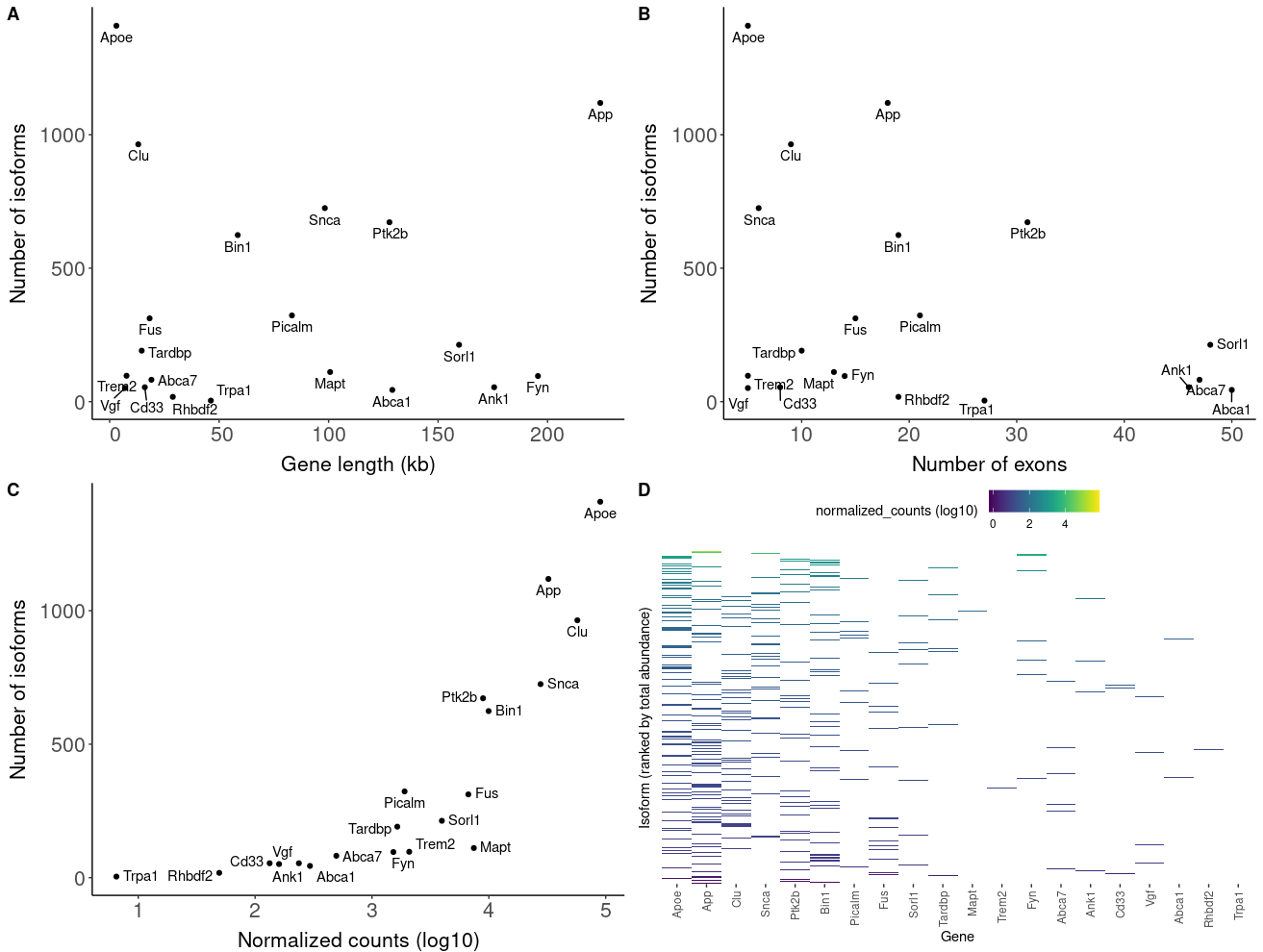

**Supplementary Figure 6: Usage of alternative splice sites and exon skipping events are commonly observed in genes associated with AD.**

Shown are bar-plots of **(A)** the number of isoforms classified with alternative 5’ and 3’ splice sites, and **(B)** the number of isoforms with exon-skipping events. Bar-plots were generated from *FICLE.* Of note, isoforms can be classified multiple times with different alternative splice sites. Also shown are examples of **(C)** known (FSM, ISM) and novel (NNC) isoforms annotated to *App* and **(D)** *Bin1*, characterized by no (green) and extensive (red) exon skipping events. Examples were found using *FICLE* and visualized using *ggtranscript*.

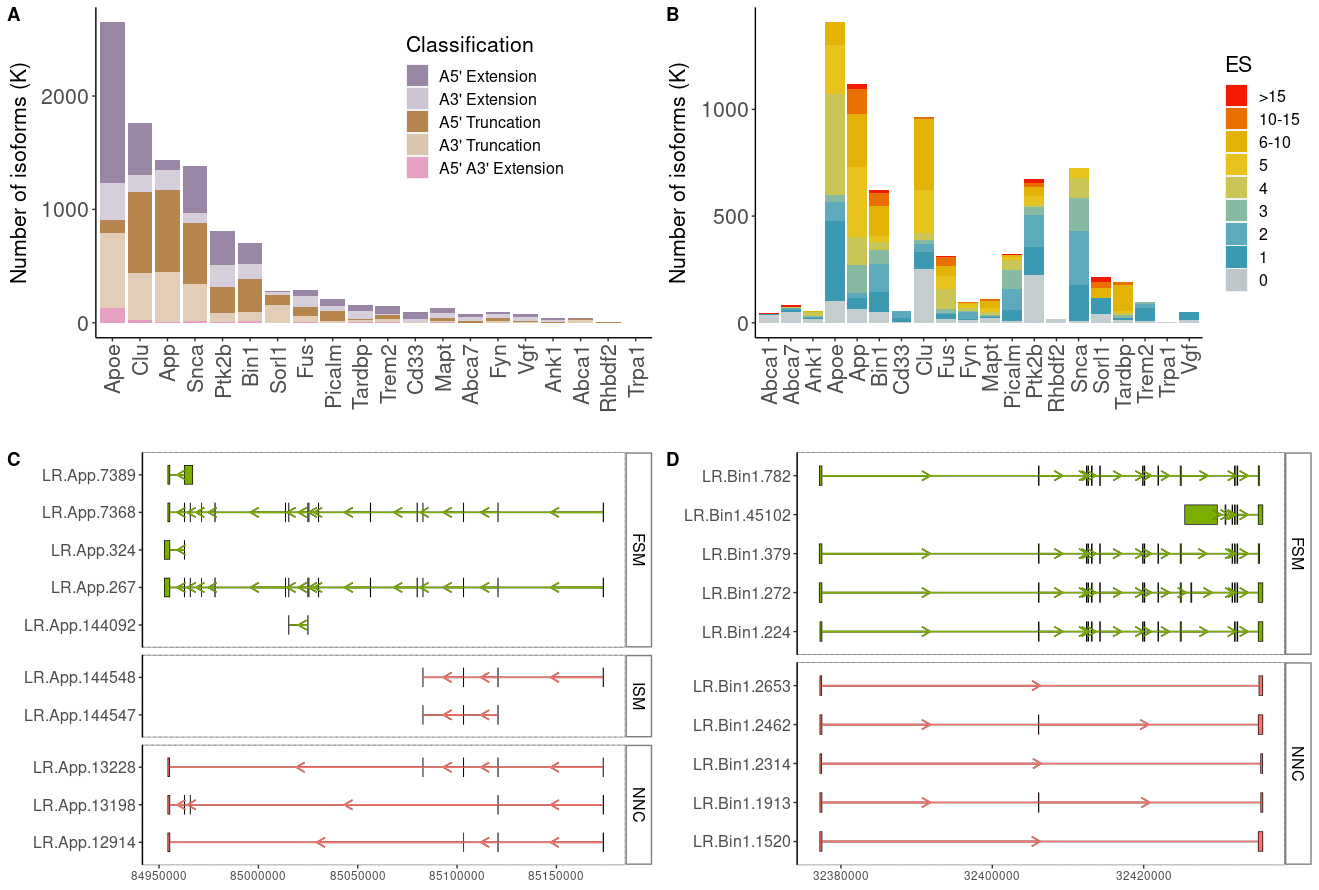

**Supplementary Figure 7: Intron retention events, although less frequently detected than exon skipping, are also observed in genes associated with AD.**

Shown are **(A)** reference *Cd33* transcripts and a novel isoform characterized by two distinct intron retention events, identified using *FICLE.* **(B)** Bar-plot of the number of isoforms with intron retention (IR) events, and the **(C)** number of isoforms with IR events spanning across multiple exons. Plots were generated using *FICLE*. **(D)** A box-plot of the expression of transcripts with 0, 1 and 2 intron retention events. **(E)** Visualization of reference *Tardbp* transcripts and isoforms with IR events spanning across multiple exons, noting the complexity of the 3’end of the gene body. Examples were found using *FICLE* and visualized using *ggtranscript*.
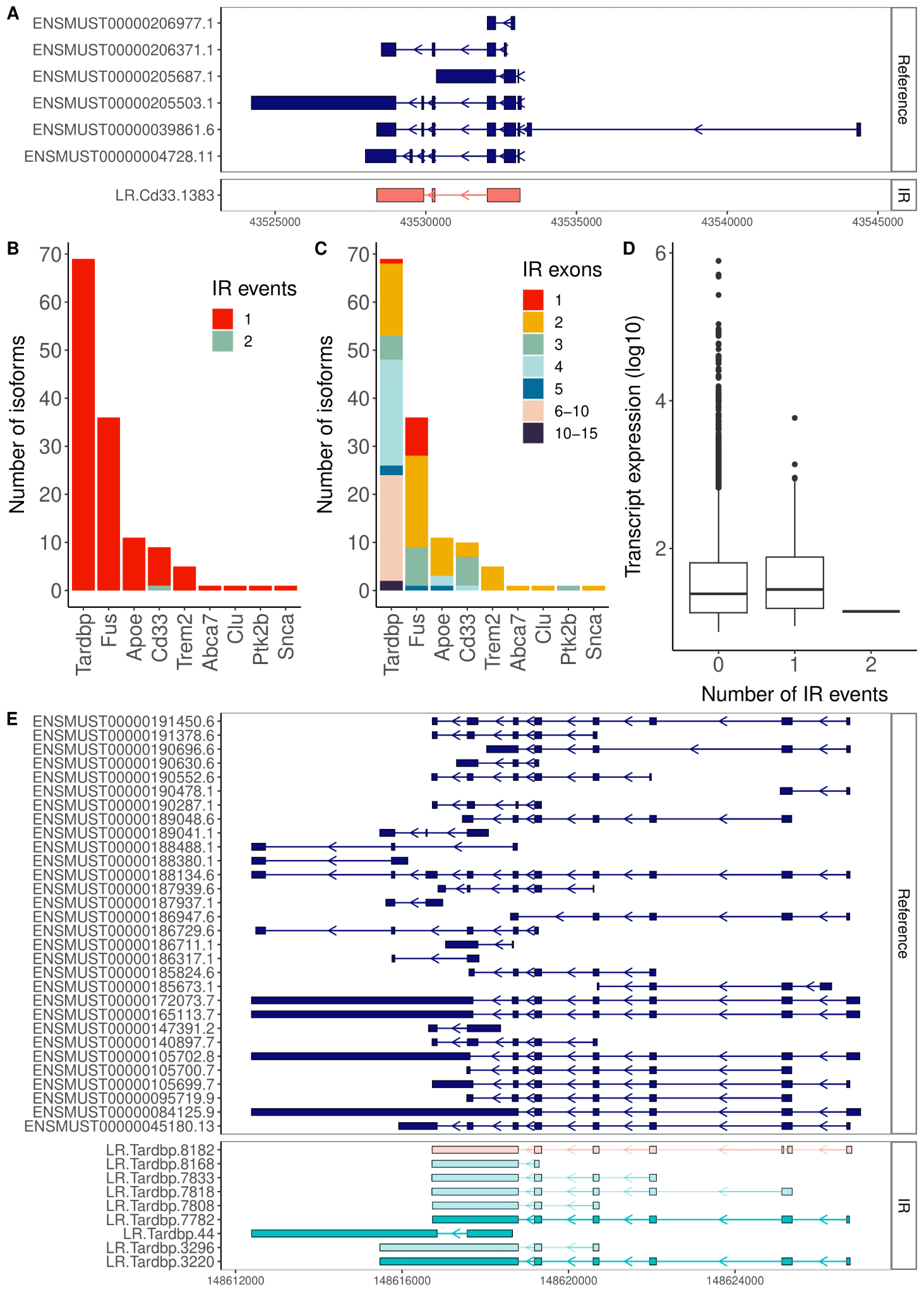

**Supplementary Figure 8: Top-ranked differentially expressed transcripts between WT and TG mice.**

Shown are box-plots of the **(A - J)** transcript expression of the 10 top-ranked differentially expressed transcripts between WT and TG mice (genotype). Transcript expression is determined from normalized ONT full length read count (Targeted ONT dataset). Differential transcript expression is performed using the Wald test in *DESeq2* (~ genotype). *SQANTI3* structural categories (FSM – Full Splice Match, ISM – Incomplete Splice Match, NIC – Novel in Catalog, NNC – Novel Not in Catalog) are provided in parenthesis for each transcript. WT – wild type, TG – transgenic.

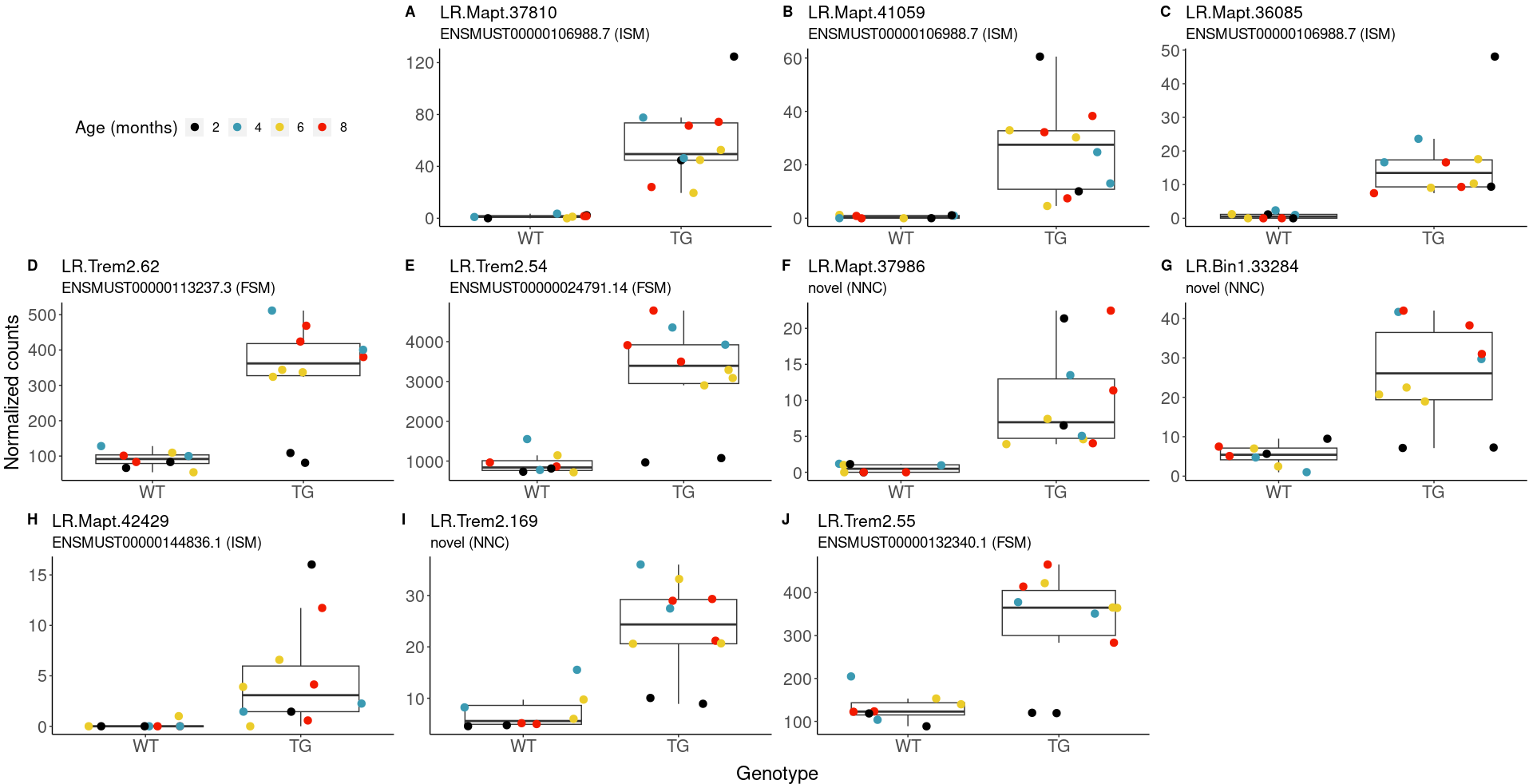

**Supplementary Figure 9: Differential transcript usage**

Shown are box plots of the transcript usage (isoform fraction) of the genes that were significantly altered **(A - C)** between WT (gray) and TG (black) mice and **(D - F)** associated with the progression of tau pathology in TG (black) mice. The usage of the top 3 most abundant transcripts are shown for each gene.

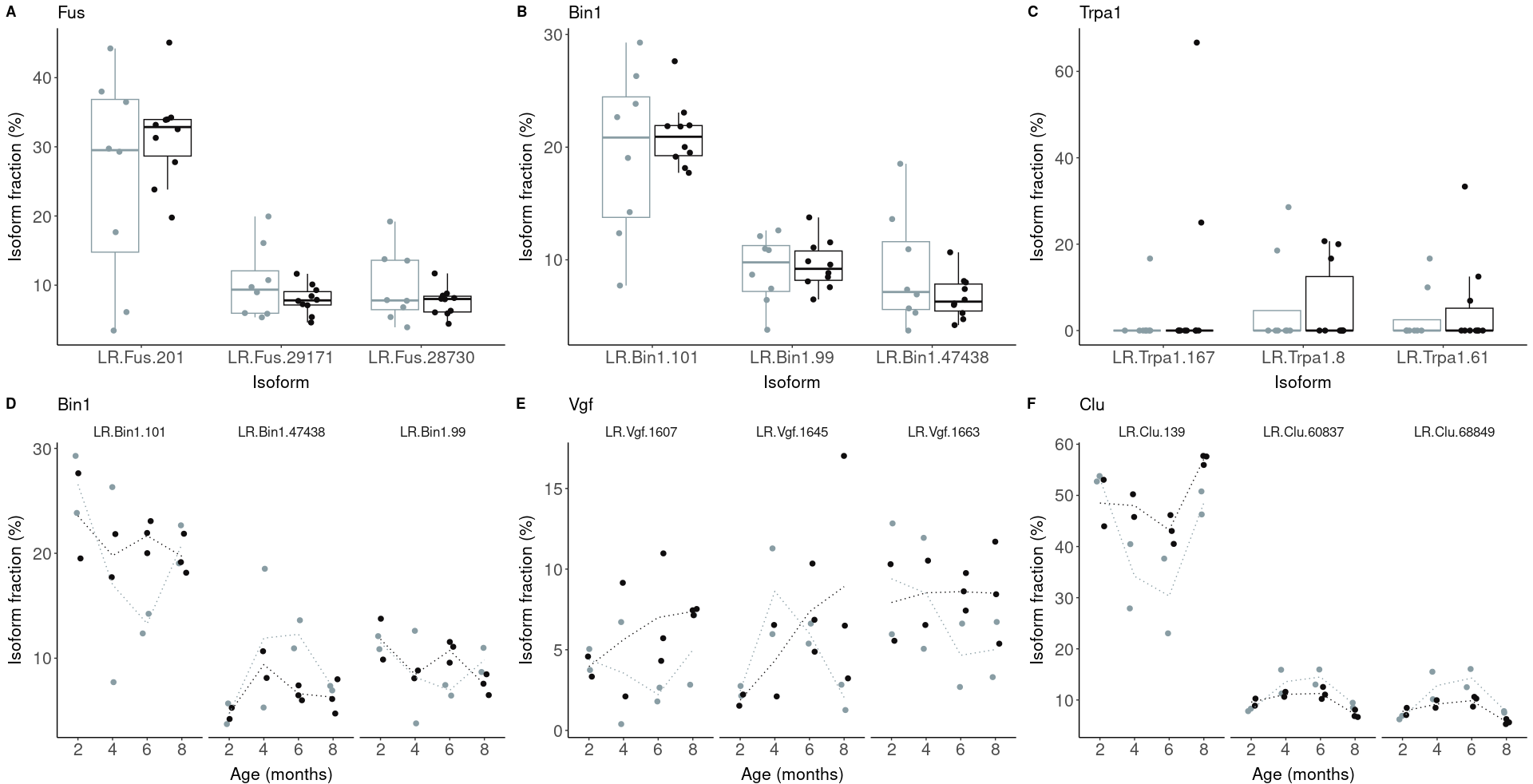

**Supplementary Figure 10: Further characterization of *Trem2* in rTg4510 mouse model**

**(A)** UCSC track of RNA-Seq data generated from matched samples, mouse reference transcripts (mm10, GENCODE) and Pfam domains of *Trem2*. The figure shows the variability of exon 2 (highlighted in red) - which encodes the V-set domain - confirmed by RNA-Seq data. **(B)** UCSC track of i) a selection of *Trem2*-associated isoforms, detected from targeted sequencing of rTg4510 mice, characterized with novel exons upstream of (highlighted in green) and within (highlighted in orange) the gene body, ii) the open reading frame (ORF; start is highlighted in blue) of shown *Trem2*-associated isoforms, showing that the novel upstream exon does not encode novel start exons whereas the internal novel exons (highlighted in orange) are retained within the ORF.

**(A)**

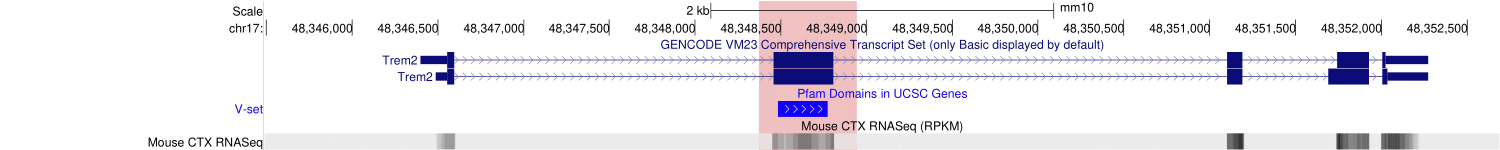

**(B)**

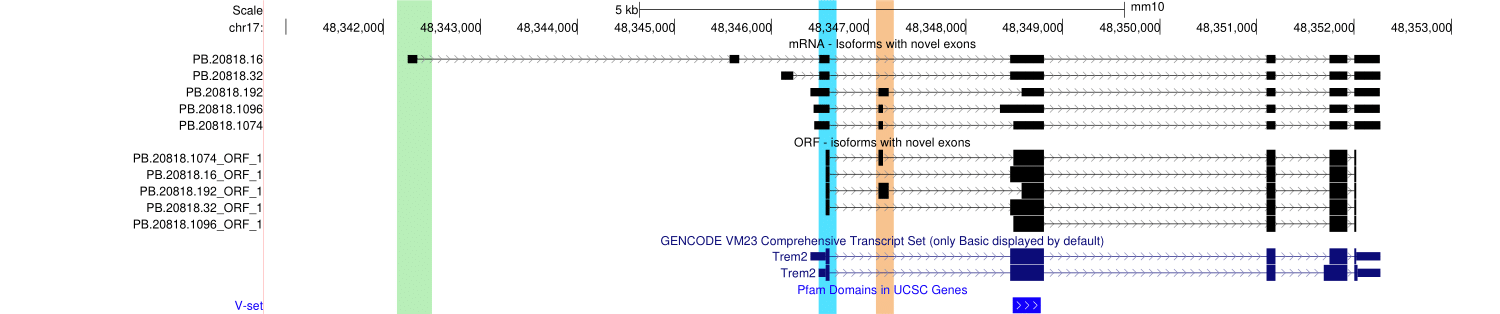

**Supplementary Figure 11: High homology between mouse and human *Bin1/BIN1* isoforms**

**(A)** NCBI BLAST [output](https://www.ncbi.nlm.nih.gov/projects/sviewer/?id=lcl%7CQuery_7439&tracks=%5Bkey:sequence_track,name:Sequence,display_name:Sequence,id:STD649220238,annots:Sequence,ShowLabel:true,ColorGaps:false,shown:true,order:1%5D%5Bkey:alignment_track,name:U762JAA0FC121,display_name:(U)%20BLAST%20Results%20for%5C:%20ENST00000316724.10%5C%7CENSG00000136717.15%5C%7COTTHUMG00000131465.3%5C%7COTTHUMT00000254298.3%5C%7CBIN1-202%5C%7CBIN1%5C%7C2487%5C%7Cprotein_coding%5C%7C,id:U762JAA0FC121,data_key:y3Rdpat2fFdQWEpQe0l0XicjJSItIAEqCTIHGJIUW7xhihBc6gDbPH6dSOQd-ETgVsgByULXUsZI3FjDbu5X7HfcUQ,dbname:NetCache,annots:BLAST%20Results%20for%5C:%20ENST00000316724.10%5C%7CENSG00000136717.15%5C%7COTTHUMG00000131465.3%5C%7COTTHUMT00000254298.3%5C%7CBIN1-202%5C%7CBIN1%5C%7C2487%5C%7Cprotein_coding%5C%7C_UUD1688391259DUU_nucleotide,Layout:Adaptive,StatDisplay:15,Color:Show%20Differences,UnalignedTailsMode:glyph,sort_by:,LinkMatePairAligns:false,ShowAlnStat:false,AlignedSeqFeats:false,Label:true,IdenticalBases:false,shown:true,order:4%5D&key=Mo2kXFKPha6pobOpgrCNp97a3NvU2fjT8Mv-4WvtokWYc3PEVhlnNUR5VQAAHFkESywcLV8zTyJVOEUncwpKCGo4TA&v=214:2362&c=993366&select=null&slim=0) of ENST00000316724.10 (human *BIN1* isoform 1) as subject and LR.Bin1.99 (ENSMUST00000025239.8, mouse *Bin1*-201 isoform) as query. **(B)** NCBI BLASTt [output](https://www.ncbi.nlm.nih.gov/projects/sviewer/?id=lcl%7CQuery_7439&tracks=%5Bkey:sequence_track,name:Sequence,display_name:Sequence,id:STD649220238,annots:Sequence,ShowLabel:true,ColorGaps:false,shown:true,order:1%5D%5Bkey:alignment_track,name:U762JAA0FC121,display_name:(U)%20BLAST%20Results%20for%5C:%20ENST00000316724.10%5C%7CENSG00000136717.15%5C%7COTTHUMG00000131465.3%5C%7COTTHUMT00000254298.3%5C%7CBIN1-202%5C%7CBIN1%5C%7C2487%5C%7Cprotein_coding%5C%7C,id:U762JAA0FC121,data_key:y3Rdpat2fFdQWEpQe0l0XicjJSItIAEqCTIHGJIUW7xhihBc6gDbPH6dSOQd-ETgVsgByULXUsZI3FjDbu5X7HfcUQ,dbname:NetCache,annots:BLAST%20Results%20for%5C:%20ENST00000316724.10%5C%7CENSG00000136717.15%5C%7COTTHUMG00000131465.3%5C%7COTTHUMT00000254298.3%5C%7CBIN1-202%5C%7CBIN1%5C%7C2487%5C%7Cprotein_coding%5C%7C_UUD1688391259DUU_nucleotide,Layout:Adaptive,StatDisplay:15,Color:Show%20Differences,UnalignedTailsMode:glyph,sort_by:,LinkMatePairAligns:false,ShowAlnStat:false,AlignedSeqFeats:false,Label:true,IdenticalBases:false,shown:true,order:4%5D&key=Mo2kXFKPha6pobOpgrCNp97a3NvU2fjT8Mv-4WvtokWYc3PEVhlnNUR5VQAAHFkESywcLV8zTyJVOEUncwpKCGo4TA&v=214:2362&c=993366&select=null&slim=0) of ENST00000409400.1 (human *BIN1* isoform 9) as subject and LR.Bin1.224 (ENSMUST00000234496.1, mouse *Bin1*-205 isoform) as query.

**(A)**
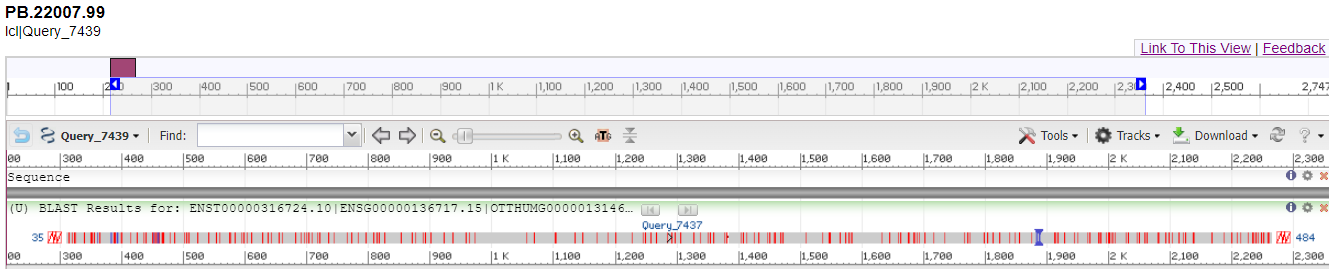

**(B)**

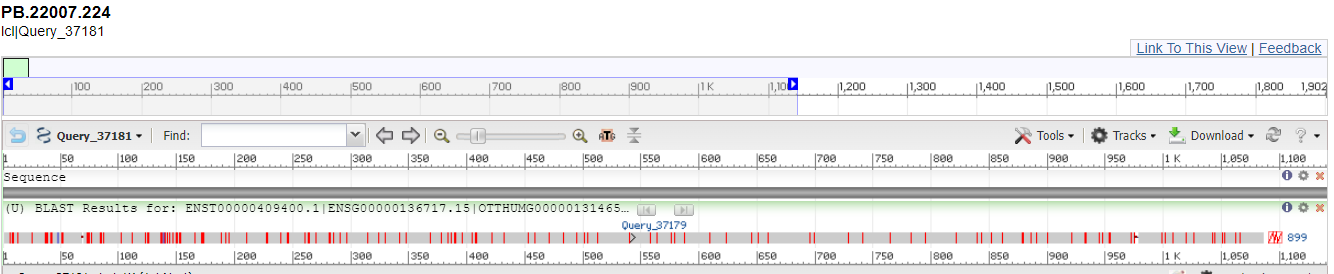
