## Supplementary Figure 1-4 for "Long-read transcript sequencing identifies differential isoform expression in the entorhinal cortex in a transgenic model of tau pathology"

**Supplementary Tables**

| Supplementary Table 1 | Sample information |
| --- | --- |
| Supplementary Table 2 | Isoform diversity in whole transcriptome dataset |
| Supplementary Table 3 | Differential gene expression analysis in whole transcriptome |
| Supplementary Table 4 | Differential transcript expression (DTE) analysis in whole transcriptome |
| Supplementary Table 5 | DTE analysis (pathology) in targeted transcriptome |
| Supplementary Table 6  Supplementary Table 7 | DTE analysis (genotype) in targeted transcriptome  Differential transcript usage analysis in targeted transcriptome |

**Supplementary Table 1. Detailed information for the samples sequenced in this study.**
“X” denotes the samples that were sequenced per dataset. TG – rTg4510 transgenic mice, WT – Wild-type mice. * Long-read RNA-Seq data (whole transcriptome and targeted profiling) generated in this study. + Short-read RNA-Seq data from Castanho *et al.* (2020).

| Sample | Genotype | Age (months) | Whole transcriptome  PacBio* | Targeted  PacBio* | Targeted  ONT* | RNA-Seq Illumina^+^ |
| --- | --- | --- | --- | --- | --- | --- |
| 1 | WT | 2 | X | X |  | X |
| 2 | WT | 2 | X | X | X | X |
| 3 | WT | 2 | X | X | X | X |
| 4 | WT | 4 |  | X |  | X |
| 5 | WT | 4 |  | X | X | X |
| 6 | WT | 4 |  | X | X | X |
| 7 | WT | 6 |  | X |  | X |
| 8 | WT | 6 |  | X | X | X |
| 9 | WT | 6 |  | X | X | X |
| 10 | WT | 8 | X | X |  | X |
| 11 | WT | 8 | X | X | X | X |
| 12 | WT | 8 | X | X | X | X |
| 13 | TG | 2 | X | X |  | X |
| 14 | TG | 2 | X | X | X | X |
| 15 | TG | 2 | X | X | X | X |
| 16 | TG | 4 |  | X |  | X |
| 17 | TG | 4 |  | X | X | X |
| 18 | TG | 4 |  | X | X | X |
| 19 | TG | 6 |  | X | X | X |
| 20 | TG | 6 |  | X | X | X |
| 21 | TG | 6 |  | X | X | X |
| 22 | TG | 8 | X | X | X | X |
| 23 | TG | 8 | X | X | X | X |
| 24 | TG | 8 | X | X | X | X |

**Supplementary Table 2. Isoform diversity in rTg4510 mouse model between genotype and age groups from whole transcriptome PacBio Iso-Seq dataset.** FSM – Full Splice Match, ISM – Incomplete Splice Match, NIC – Novel in Catalog, NNC – Novel Not in Catalog.

|  | Wild-type mice | | | rTg4510 transgenic mice | | |
| --- | --- | --- | --- | --- | --- | --- |
|  | 2 & 8 months | 2 months | 8 months | 2 & 8 months | 2 months | 8 months |
| Total number of genes | 14118 | 13191 | 13312 | 14213 | 12985 | 13616 |
| Annotated genes | 13932 (98.68%) | 13081 (99.17%) | 13168 (98.92%) | 14031 (98.72%) | 12874 (99.15%) | 13474 (98.96%) |
| Novel genes | 186 (1.32%) | 110 (0.83%) | 144 (1.08%) | 182 (1.28%) | 111 (0.85%) | 142 (1.04%) |
| Number of isoforms | 62533 | 48516 | 50278 | 63038 | 45903 | 52730 |
| FSM | 33239 (53.15%) | 27878 (57.46%) | 28689 (57.06%) | 33563 (53.24%) | 26825 (58.44%) | 29916 (56.73%) |
| ISM | 4927 (7.88%) | 3426 (7.06%) | 3841 (7.64%) | 4864 (7.72%) | 3279 (7.14%) | 3764 (7.14%) |
| NIC | 15305 (24.48%) | 11012 (22.7%) | 11407 (22.69%) | 15595 (24.74%) | 10214 (22.25%) | 12369 (23.46%) |
| NNC | 8518 (13.62%) | 5838 (12.03%) | 5953 (11.84%) | 8484 (13.46%) | 5259 (11.46%) | 6282 (11.91%) |
| Genic genomic | 63 (0.1%) | 44 (0.09%) | 44 (0.09%) | 61 (0.1%) | 32 (0.07%) | 47 (0.09%) |
| Antisense | 97 (0.16%) | 52 (0.11%) | 77 (0.15%) | 104 (0.16%) | 68 (0.15%) | 75 (0.14%) |
| Fusion | 276 (0.44%) | 200 (0.41%) | 186 (0.37%) | 268 (0.43%) | 167 (0.36%) | 196 (0.37%) |
| Intergenic | 108 (0.17%) | 66 (0.14%) | 81 (0.16%) | 99 (0.16%) | 59 (0.13%) | 81 (0.15%) |
| Genic intron | 0 (0%) | 0 (0%) | 0 (0%) | 0 (0%) | 0 (0%) | 0 (0%) |
| Isoform length (bp)  median (range) | 2691  (82 - 15016) | 2740  (88 - 15016) | 2614  (82 - 14850) | 2698  (82 - 15913) | 2548  (88 - 14302) | 2754  (82 - 15913) |
| Number of exons  median (range) | 8 (1 - 89) | 8 (1 - 89) | 8 (1 - 89) | 8 (1 - 89) | 8 (1 - 77) | 9 (1 - 89) |
| Number of isoforms with 50 bp CAGE | 52096 (83.31%) | 40589 (83.66%) | 42378 (84.29%) | 52633 (83.49%) | 38227 (83.28%) | 44729 (84.83%) |

**Supplementary Table 3. Differentially expressed genes associated with the progression of tau pathology in the whole transcriptome PacBio Iso-Seq dataset.**

Gene expression is determined from the summation of normalized Iso-Seq full-length read counts (whole transcriptome dataset) of associated transcripts. Differential gene expression is performed using the Wald test in *DESeq2* (~ genotype + age + genotype * age). The counts of the genotype groups refer to the mean normalized full-length reads. TG – rTg4510 transgenic mice, WT – Wild-type mice.

| Associated gene | log2FoldChange | lfcSE | P-value | FDR | WT 2 months  counts | WT 8 months  counts | TG 2 months  counts | TG 8 months  counts |
| --- | --- | --- | --- | --- | --- | --- | --- | --- |
| Gfap | 3.37 | 5.96E-01 | 1.61E-08 | 1.99E-04 | 83.2 | 70.1 | 117 | 1020 |
| C4b | 5.01 | 9.32E-01 | 7.79E-08 | 4.83E-04 | 4.66 | 2.47 | 6 | 104 |

**Supplementary Table 4. Differentially expressed transcripts associated with the progression of tau pathology in the whole transcriptome PacBio Iso-Seq dataset.**

Transcript expression is derived from normalized Iso-Seq full-length read counts (whole transcriptome dataset). Differential transcript expression is performed using the Wald test in *DESeq2* (~ genotype + age + genotype * age). log2FC – log_2_ fold-change, mos – months, TG – transgenic, WT – wild-type.

| Associated gene | Isoform | Associated transcript | log2FC | lfcSE | P-value | FDR | TG 2mos counts | TG 8mos counts | WT 2mos counts | WT 8mos counts |
| --- | --- | --- | --- | --- | --- | --- | --- | --- | --- | --- |
| Gfap | PB.2973.16 | ENSMUST00000067444.9 | 3.57 | 6.44E-01 | 2.95E-08 | 1.16E-03 | 89.1 | 849 | 68 | 54.5 |
| C4b | PB.7022.9 | ENSMUST00000069507.8 | 5.51 | 1.18E+00 | 3.26E-06 | 6.39E-2 | 3.69 | 76.7 | 3.31 | 1.59 |

**Supplementary Table 5. Differentially expressed transcripts between WT and rTg4510 mice across age (pathology) in targeted ONT and Iso-Seq datasets.**

See attached xlsx file SupplementaryTable5. Transcript expression is derived from normalized ONT and Iso-Seq full-length read counts (ONT and Iso-Seq targeted datasets). Differential transcript expression was performed using the Wald test in *DESeq2* (~ genotype + age + genotype * age) across all ages in ONT and Iso-Seq datasets. *SQANTI3* structural categories (FSM – Full Splice Match, ISM – Incomplete Splice Match, NIC – Novel in Catalog, NNC – Novel Not in Catalog) and subcategories are provided in parenthesis for each transcript. WT – wild type, TG – transgenic. Significant transcripts (ONT FDR < 0.05) are highlighted in green.

**Supplementary Table 6.** Differentially expressed transcripts between WT and rTg4510 mice (genotype) in the targeted ONT and Iso-Seq datasets.

See attached xlsx file SupplementaryTable6. Transcript expression is derived from the normalised ONT and Iso-Seq full-length read counts (ONT and Iso-Seq targeted datasets). Differential transcript expression was performed using the Wald test in *DESeq2* (~ genotype) between WT and TG mice in ONT and Iso-Seq datasets. *SQANTI3* structural categories (FSM – Full Splice Match, ISM – Incomplete Splice Match, NIC – Novel in Catalog, NNC – Novel Not in Catalog) and subcategories are provided in parenthesis for each transcript. WT – wild type, TG – transgenic. Significant transcripts (ONT FDR < 0.05) are highlighted in green.

**Supplementary Table 7. Genes with differential transcript usage between WT and rTg4510 mice in targeted dataset.**

See attached xlsx file SupplementaryTable7. Transcript expression is derived from the normalized ONT full-length read counts (ONT targeted datasets). Differential transcript usage between experimental groups (Genotype: WT and TG, Pathology: WT and TG across 4 ages) was performed using *EdgeR spliceVariants* in *tappAS* with filtering of minor isoforms using fold-change (log2) (see **Methods**). A “TRUE” podium change indicates a switch in major isoform usage.
